# CGRP receptor-expressing neurons in the central amygdala contributes to injury-induced pain hypersensitivity

**DOI:** 10.64898/2026.04.02.716115

**Authors:** Sudhuman Singh, Ana Danko, Benjamin Neugebauer, Sarah Chaudhry, Lakeisha A Lewter, William Fortún, Jenny Lin, Spring Valdivia, Torri D. Wilson, Jeitzel M. Torres-Rodriguez, Benedict J Kolber, Yarimar Carrasquillo

## Abstract

The central nucleus of amygdala (CeA) comprises diverse populations of neurons, forming a complex network responsible for regulating various behavioral responses. Among these, neurons expressing calcitonin gene-related peptide receptors (CGRPR) have emerged as key players in CGRP neuropeptide-mediated pain modulation. While previous studies emphasize CGRP’s key role in synaptic plasticity and its connection with pain behavior in the CeA, the precise functional attributes and contributions of CeA-CGRPR-expressing neurons in pain processing remain elusive. This study reveals the co-localization of CGRPR-expressing neurons in the CeA with phosphorylated extracellular signal-regulated kinase (pERK), a marker indicating pain plasticity, in a neuropathic pain model. Electrophysiological assessments of these neurons in slice preparations unveiled heightened intrinsic excitability after sciatic nerve cuff implantation, contingent upon their rostro-caudal positioning within the CeA. Furthermore, our behavioral experiments using chemogenetic inhibition of CeA-CGRPR neurons demonstrated the ability to reverse nerve injury-induced hypersensitivity. Conversely, activating these neurons induced pain-related hypersensitivity even in the absence of injury. Our findings also highlight a sex-specific role of CeA-CGRPR neurons in formalin-induced spontaneous pain response. Collectively, these data reinforce the involvement of CeA-CGRPR neurons in pain processing, contributing to a better understanding of how neural circuits are affected in persistent pain conditions.

**Significance Statement:** Our study shows the role of CGRPR-expressing neurons within the CeA during pain processing. Using a cuff-implanted neuropathic mouse model, we discovered that CGRPR-expressing neurons co-localize with phosphorylated extracellular signal-regulated kinase (pERK), a hallmark of pain plasticity, in both male and female mice. Furthermore, our electrophysiological investigations reveal that posterior CeA-CGRPR neurons exhibit increased excitability following sciatic nerve cuff implantation. Importantly, we demonstrate that CeA-CGRPR neurons exert bidirectional effects on pain behavior in mice, irrespective of sex differences in nerve injury-induced pain responses while showing sex-specific spontaneous pain responses in the formalin-induced model. These findings show the role of CeA-CGRPR neurons in pain modulation, underscoring their potential significance in understanding and addressing persistent pain conditions.

## Introduction

The central amygdala (CeA) is a complex brain region within the limbic system that plays a critical role in emotional processing, fear, anxiety, and pain modulation (Ehrlich et al., 2009; Li et al., 2013; Penzo et al., 2015; Moscarello and Penzo 2022; Calhoon and Tye, 2015; Ahrens et al., 2018; Bernard et al., 1992; Wilson et al., 2019; Veinante et al., 2013; Singh et al., 2022). It is composed of distinct populations of neurons exhibiting remarkable heterogeneity, forming a complex network of functionally and genetically determined cell types responsible for regulating various behavioral responses (Janak and Tye 2015; O’ Leary et al., 2022; Fadok et al., 2018). Recent studies have demonstrated a dual role of the CeA in nociception, driving either pronociceptive or antinociceptive based on the specific cell types involved (Wilson et al., 2019). These results suggest that the CeA functions as a rheostat of pain modulation. These findings reinforce the importance of unraveling the precise mechanisms underlying the interplay between CeA circuits and pain regulation. Within CeA, CGRPR (calcitonin gene-related peptide receptor; calcitonin receptor-like receptor: *Calcrl*) expressing neurons garnered attention due to their involvement in CGRP-mediated pain modulation (Xu et al., 2003; Han et al., 2005, 2010; Allen et al., 2023). CGRP is a neuropeptide comprised of 37 amino acids, produced by neurons in both the peripheral and central nervous systems (Russo, 2015). It functions as a potent vasodilator and plays a role in transmitting pain signals (Iyengar et., 2017; Neugebauer., 2022). The main source of CGRP in the amygdala comes from CGRP-containing neurons in lateral parabrachial nucleus (LPB) through the spino-parabrachio-amygdaloid pain pathway (Dobolyi et al., 2005; D’Hanis., 2007; Han et al., 2015). Studies using genetic labeling techniques have shown that CGRP receptors are primarily expressed in the lateral and capsular divisions of the amygdala, with a higher level of expression in PKCδ neurons (Han et al., 2015). Many studies have shown that CeA-CGRP receptors are targeted by CGRP and play a significant role in pain-related synaptic plasticity (Han et al., 2005; Shinohara et al., 2017). Studies conducted by Han et al. (2005) and Shinohara et al. (2017) have demonstrated CGRP’s influential role in modulating synaptic input from the PB to the CeA. These investigations demonstrate the significance of CGRP in shaping the synaptic connections within the CeA and its key involvement in pain-related neural circuitry. However, the specific functional properties and contributions of CGRPR-expressing neurons in CeA in pain processing are still the subject of active investigation.

Based upon prior research defining the heterogeneous nature of neuronal subtypes within the CeA and role of CGRP peptide in synaptic plasticity and pain regulation, our hypothesis centered around the potential contribution of CeA-CGRPR cells in pain processes. Using a combined approach of cellular electrophysiological, and *in vivo* chemogenetics, we aimed to dissect the specific contributions of these neurons to pain modulation and synaptic plasticity. We found that nerve injury triggers phosphorylated extracellular signal-regulated kinase (pERK) expression in CeA-CGRPR neurons, indicating their involvement in pain plasticity (Carrasquillo and Gereau., 2007). Moreover, the increased excitability of CGRPR-expressing neurons during pain depends on their rostral-caudal level within the CeA. Chemogenetic inhibition of these neurons effectively reverses hypersensitivity caused by cuff-induced nerve injury, while activating them alone induces pain-related hypersensitivity even without injury. Additionally, our findings revealed sex-specific modulation in the formalin-induced spontaneous pain response, highlighting the involvement of CeA-CGRPR neurons in pain regulation.

## Materials and Methods

### Animals

Adult male and female CGRPR-cre mice, aged 8-17 weeks, were used in this study. The CGRPR-cre mice, kindly provided by Dr. Richard Palmiter (University of Washington), were previously generated and their generation has been described by Han, Soleiman et al. 2015. These mice are also available through The Jackson Laboratory (Strain #:041390 B6.Cg-*Calcrl^tm1.1(cre/EGFP)Rpa^*/J) and contain a cre-recombinase and GFP transgene downstream of the normal *Calcrl* gene. To generate CGRPR-Ai9 reporter mice, homozygous CGRPR-cre mice were backcrossed with homozygous Ai9B6J mice (The Jackson Laboratory, Strain #:007909 B6.Cg-*Gt(ROSA)26Sor^tm9(CAG-tdTomato)Hze^*/J). Offspring were genotyped using tail biopsies for DNA extraction and subjected to PCR (Transnetyx) using the following primers: CAGGCTAAGTGCCTTCTCTACA (reverse) and TTAATCCATATTGGCAGAACGAAAACG (forward) to confirm the presence of cre-recombinase. The mice were housed in a vivarium under control conditions (temperature: 22 ± 2°C, humidity: 50 ± 10 %) with a reversed 12-hour light/dark cycle (9 PM to 9 AM light) and provided with *ad libitum* access to food and water. All behavioral tests were conducted during the dark period using red light illumination. Mice received daily intraperitoneal injections (i.p.) of 100 µl of saline and were habituated by the experimenter for one week prior to the commencement of behavioral and electrophysiological experiments, following the cupping method as described by Hurst and West (2010). Following surgeries, littermates with the same treatment were co-housed in divider cages (up to a maximum of 2 mice per cage, one in each compartment). All procedures strictly followed the guidelines set forth by the National Institutes of Health (NIH) and were approved by the Animal Care and Use Committee of the National Institute of Neurological Disorders and Stroke and the National Institute of Deafness.

### Stereotaxic surgery

Mice were initially anesthetized with 5% isoflurane to prepare for the surgery. Once anesthetized, the mice were securely positioned in a head-fixed manner on a stereotaxic apparatus (David Kopf Instruments), and the anesthesia level was maintained at 1.5-2 % isoflurane with a flow rate of 0.5 L/min throughout the surgery. To ensure thermal stability, a hand warmer was utilized during the procedure. For the fidelity experiment, the right central amygdala (CeA) of CGRPR-Cre mice were infected with 0.05 µl of AAV8-hSyn-DIO-mCherry (Addgene viral prep #50459-AAV8), or AAV8-hSyn-DIO-EGFP (UNC vector core, Chapel Hill, NC). For the chemogenetic experiment, 0.05 µl of AAV8-hSyn-DIO-mCherry (Addgene viral prep #50459-AAV8), or AAV8-hSyn-DIO-hM3Di-mCherry (Addgene viral prep #44362-AAV8), or AAV8-hSyn-DIO-hM3Dq-mCherry (Addgene viral prep #44361-AAV8) virus was microinjected into the right CeA using a 32-gauge, 0.5 μl Hamilton Neuros syringe. The injections were performed at a flow rate of 0.1 μl/min, and the injector was left in place for an additional 5 minutes at the end of the injection to allow for virus diffusion. The following stereotaxic coordinates were used to target the CeA in CGRPR-cre mice: 1.25 mm posterior from bregma, 3.05 mm lateral to the midline, 4.55 mm ventral to the skull surface. Mice were allowed to recover for 1 week before additional experimental procedures were conducted. At the end of each experiment, mice were transcardially perfused with a solution of 4 % paraformaldehyde in 0.1 M Phosphate Buffer (PFA/PB), pH 7.4, and their brains were stained for mCherry as described below to verify the injection site. The anatomical boundaries of each region were identified using a mouse brain atlas (Paxinos and Franklin, 2001), and drawings illustrating the spread of the virus at different rostro-caudal levels were created for each infected mouse. Only mice that exhibited virus injections restricted to the CeA were included in the behavioral experiments.

### Evaluation of CeA-Calcrl-Cre expression by RNAScope

To determine the validity and efficiency of Calcrl-Cre expression, we imaged *Calcrl* mRNA colocalized with fluorescent reporters dependent on the Cre-recombinase. This included (1) the expression of viral-mediated fluorescent reporters infected into adult animals (brains from *Calcrl*-Cre animals infected with AAV-DIO-mCherry or AAV-DIO-GFP) and (2) the expression of developmentally expressed tdTomato fluorescent reporters from double transgenic mice (CalcrlCre:Ai14, and CalcrlCre:Ai9 animals). All brains were frozen, cryoprotected, and sectioned at 20 μm. *In situ* hybridization RNAscope was performed on representative CeA sections using the RNAscope probe for *Calcrl (refs:*C1-C3 452281). Following RNAScope, immunohistochemistry (IHC) was performed to identify virus-labeled or reporter-labeled cells. The following antibodies were used to perform GFP, mCherry, and Tdtomato IHC, respectively: (Aves - GFP-1010, 1:500; Abcam - ab167453, 1:500; Sicgen - AB8181, 1:500). Slides were imaged on an Olympus IX73 at 20 X magnification. Only animals with accurate targeting for the CeA were included in the analysis. Cells were considered positive for the fluorescent marker if they colocalized with DAPI. Positive *Calcrl* positive cells were identified as a DAPI-labeled nucleus surrounded by at least 3 fluorescent puncta indicative of positive *Calcrl* mRNA. Cell percent colocalization was averaged across sections from 3 different animals for virus-infected and transgenic animals. Cre fidelity was quantified by *Calcrl* positive mRNA divided by fluorescent marker multiplied by 100 %. This quantifies if the reporters are truly expressed in *Calcrl-*positive cells. Efficiency was quantified by fluorescent marker divided by *Calcrl* positive mRNA multiplied by 100 %. This quantifies how much of the endogenous *Calcrl* positive population is targeted by the virus or developmental tdTomato expression.

### Sciatic cuff surgery

The sciatic cuff implantation procedure was conducted on mice one week after the brain surgeries, following the methodology outlined in Benbouzid et al. (2008). The mice were initially anesthetized with 2 % isoflurane at a flow rate of 0.5 L/min. An incision of approximately 1 cm was made in the proximal one-third of the lateral left thigh, and the sciatic nerve was exposed and gently stretched using forceps. In the group receiving the cuff implant, a 2 mm-long polyethylene tubing (PE 20) with a 0.38 mm inner diameter and 1.09 mm outer diameter was carefully placed around the sciatic nerve and then returned to its original position. The sham group underwent a similar procedure of sciatic nerve exposure and stretching but without any tubing implantation. Following the procedure, the incision on the thigh was closed with wound clips, and the mice were given at least one week to recover before conducting any follow-up experiments.

### *Ex vivo* electrophysiology

#### Acute slice preparation

*Ex vivo* slice electrophysiology experiments were conducted 7-15 days after sciatic nerve surgery. On the day of the experiment, following an i.p. injection of avertin anesthesia (1.25 %, 0.025 mL/g body weight), mice were fastened to a dissection tray, the heart was exposed, and a small incision was made in the right atrium while inserting a butterfly syringe into the left ventricle. Mice were then transcardially perfused with approximately 60 ml of choline chloride-based cutting solution containing: 110.0 mM choline chloride, 12.7 mM L-ascorbic acid, 3.1 mM pyruvic acid, 25.0 mM D-Glucose, 7.2 mM MgCl_2_, 0.5 mM CaCl_2_, 25.0 mM NaHCO_3_, 1.25 mM NaH_2_PO_4_, 2.5 mM KCl. Brains were then swiftly but delicately extracted, and a tissue block containing the right amygdala was isolated using a metal blade and fastened in the collection chamber of the VT1200 S vibratome (Leica Microsystems Inc.) using Loctite 404 Quickset adhesive (Ted Pella). Ice-cold cutting solution was quickly added to the collection chamber and coronal slices, of 250 μm thickness, were collected and transferred for 30 minutes to a recovery chamber containing oxygenated artificial cerebrospinal fluid (ACSF) (25.0 mM D-glucose, 1.0 mM MgCl_2_, 2.0 mM CaCl_2_, 25.0 mM NaHCO_3_, 1.25 mM NaH_2_PO_4_, 2.5 mM KCl, 125.0 mM NaCl), heated to 33°C. After this, the recovery chamber was left at room temperature for an additional 20 minutes. Both cutting solution and ACSF were oxygenated continuously with 95 %/5 % O_2_/CO_2_.

#### Whole-cell patch-clamp electrophysiology

After recovery, slices were transferred to the heated recording chamber continuously perfused with oxygenated ACSF at a rate of 1mL/min and attached to an inline solution heater (Warner Instruments). Temperature was maintained at 33°±1°C. Patch pipettes ranged in resistance from 2 to 5 MΩ and were filled with a potassium methylsulfate internal solution (120.0 mM KMeSO_4_, 20.0 mM KCl, 10.0 mM HEPES, 0.2 mM EGTA, 8.0 mM NaCl, 4.0 mM Mg-ATP, 0.3 mM Tris-GTP, 14.0 mM phosphocreatine, solution pH adjusted to 7.3 with KOH, ∼300 mosmol^-1^). Current clamp recordings were performed in the laterocapsular subdivision of the CeA, identified using fiber bundles and anatomical landmarks as a reference as described in the mouse brain atlas (Paxinos and Franklin 2001). Cells were identified using differential interference contrast optics with infrared illumination and fluorescent microscopy (Nikon Eclipse FN-S2N, FN1). Cre-expressing neurons were identified by an expression of tdTomato fluorescence. Recordings were performed using a Multiclamp 700B patch-clamp amplifier with a Digidata 1500 acquisition system and pCLAMP 11.2 software (Molecular Devices). Before forming a gigaseal with the cell membrane, pipette tips were zeroed. Current Clamp recordings entailed injecting 500 ms depolarizing current (0-380 pA) to assess the frequency of action potentials in fluorescently labeled neurons at resting membrane potential from acute slices prepared from CGRPR-cre:Ai9 mice. Recordings were acquired at 100 kHz and lowpass filtered at 5 kHz. Pipette capacitance and series resistance were monitored in a voltage clamp by holding the cells at -70 mV, followed by ±10 mV voltage steps of 25 ms duration. Liquid junction potentials were not corrected during recordings.

#### Data analysis and statistics

Data analysis and statistics were performed and compiled using Graphpad Prism (version 9.5.1 for Windows, Graphpad Software, San Diego, California USA, www.graphpadprism.com), Clampfit 11.2 (Molecular Devices), and Microsoft Excel. At least seven mice were used as biological replicates for each treatment. CGRPR neurons were considered late firing (LF) if they had latencies higher than 100 ms (sham) or 90 ms (cuff) at a current injection step that elicited 4-5 action potentials. The range of current injections that elicited 4 to 5 action potentials in LF neurons was 120-380 pA. The majority of cells were classified as LF and therefore analyses were restricted to this firing type. Anterior cells were defined as those ranging from -0.70 to -1.46 mm relative to bregma, while posterior cells were defined as those ranging from -1.58 to -1.82 mm relative to bregma. Definitions of anterior and posterior were based on those described previously (Bowen et al., 2023). Anterior cells were defined as those collected from -0.70 to -1.46 mm (relative to bregma), while posterior cells were defined as those collected from -1.58 to -1.82 mm (relative to bregma). The excitability was plotted by tallying the number of action potentials at each respective current step and sorting the data according to sex, treatment, and rostro-caudal level, generating input/output curves as described previously (Neugebauer, Li et al. 2003, Wilson, Valdivia et al. 2019). We chose to pool male and female data as initial analyses showed no difference in sex. All results shown are expressed as mean ±SEM. Analyses were performed using student’s (unpaired) t-test or two-way analysis of variance (ANOVA) in Prism 9.5.1.

Capacitance was calculated by calculating the area (pA*ms) of capacitive transients in response to ±10 mV voltage steps of 25 ms duration. Input Resistance (Rin) was derived from the quotient of mean change in current and ±10 mV change in potential of 25 ms duration. Resting membrane potential (Vrest), was measured by finding the mean of traces in the 62.5 ms preceding a 500 ms depolarizing current injection. Rheobase was defined as the earliest depolarizing injection step which was sufficient to induce an action potential. Latency to the first spike was measured as the time (ms) between the onset of depolarizing current injection (380 pA) and the peak of the first action potential.

### Nociceptive Behaviors Assays

#### Acetone test

The acetone test is a commonly used method for assessing pain hypersensitivity (Choi et al., 1994). Briefly, mice were individually acclimated for 2-3 hours in a ventilated opaque white plexiglass testing chamber placed on an elevated platform with a wire mesh floor. A small drop of acetone was gently applied to the center of the plantar surface of the hind paw. The mice’s nociceptive responses were evaluated for 60 seconds, utilizing a modified 0-2-point system (Colburn et al., 2007). According to this scoring system, a score of 0 represented rapid, transient lifting, licking, or shaking of the hind paw, which immediately subsided; a score of 1 indicated lifting, licking, and/or shaking of the hind paw that persisted beyond the initial application but subsided within 5 seconds; a score of 2 represented protracted, repeated lifting, licking, and/or shaking of the hind paw. The test was repeated for each hind paw, with a minimum 5 minute interval between tests to allow for recovery.

#### Hargreaves test

To assess the thermal sensitivity of the paw, a modified version of the Hargreaves Method (Hargreaves et al., 1988; Wilson et al., 2019) was employed. Mice were acclimated in small plexiglass testing chambers with a glass floor for a minimum of 1-hour or until they exhibited signs of calmness. Following the habituation period, a noxious radiant light beam, generated by an apparatus (IITC Life Sciences, Woodland Hills, CA), was directed onto the midplantar surface of the hind paw through the glass floor. The latency of paw withdrawal from the radiant light beam was recorded, with a cutoff time of 15 seconds and an active intensity of 25 to prevent tissue damage. The threshold for each hind paw was determined by calculating the average of five trials conducted at 5-minute intervals.

#### von Frey filament test

To evaluate mechanical hypersensitivity, paw withdrawal responses to von Frey filaments (North Coast Medical, Inc., San Jose, CA) were measured following the protocol described by Wilson et al. (2019). Mice were acclimated on a mesh floor in individual ventilated opaque white Plexiglas chambers for a minimum of 2 hours. Once acclimated, each von Frey filament was applied to the center of the plantar surface of the hind paw for a duration of 2-3 seconds, applying enough force to cause slight bending against the paw. The smallest filament that elicited a paw withdrawal response in at least three out of five measurements was considered as the mechanical threshold for that trial. The average of five trials was calculated to determine the threshold value for each hind paw.

#### Randall-Selitto test

To assess the mechanical pressure threshold of the hind paws, the Randall-Selitto test (Randall and Selitto, 1957) was performed on lightly anesthetized mice. This test involves gradually applying increasing force to the plantar surface of the hind paw until the animal responds by withdrawing its paw or vocalizing. Initially, mice were anesthetized using 5 % isoflurane in an induction chamber. Subsequently, the animals were maintained under light anesthesia with 0.5-1 % isoflurane at a flow rate of 0.5 L/min. A maximal cut-off of 200 g force was applied to prevent any tissue damage. Five trials per animal were recorded and the average was calculated.

#### Formalin test

The formalin test was performed in both male and female CGRPR-cre mice who had received chemogenetic vectors. Acclimatized mice received an i.p. injection of either CNO (10 mg/kg) or saline on the test day. Thirty minutes after the i.p. injection, mice received 10 µL of 3 % formalin subcutaneously on the plantar surface of either their left or right hind paw. Subsequently, mice were returned to the testing chamber. Their nociceptive behaviors (including licking, lifting, and shaking of the formalin-injected paw) were observed and recorded in 5-minute bins for 40 minutes. For analysis, formalin nociceptive behavior was divided into two phases: Phase 1, 0-5 minutes post-formalin injection, and Phase 2, 5-40 minutes post-formalin injection.

### Validation of hM4Di and hM3Dq DREADDs

To confirm the functionality of the DREADDs employed in the chemogenetic behavioral assays, we evaluated the expression of the Fos protein within mCherry-expressing DREADDs neurons. We anticipated observing neuronal inhibition in hM4Di-expressing neurons and activation in hM3Dq-expressing neurons. This validation was conducted through double IHC to assess the co-expression of Fos and mCherry proteins.

### Immunohistochemistry

At the end of each experiment, mice were deeply anesthetized using 1.25% Avertin anesthesia (2,2,2-tribromoethanol and tert-amyl alcohol in 0.9 % NaCl; 0.025 ml/g body weight) administered intraperitoneally. Subsequently, the mice were transcardially perfused with 0.9 % NaCl at 37°C, followed by 100 mL of ice-cold 4 % paraformaldehyde in phosphate buffer solution (PFA/PB). The brains were dissected and post-fixed in 4 % PFA/PB overnight at 4°C. To prepare the brain tissues for immunostaining, cryoprotection was performed by immersing the tissue in 30 % sucrose/PB for 48-hours. Coronal sections with a thickness of 30 μm were obtained using a freezing sliding microtome. These sections were stored in 0.1 M Phosphate Buffered Saline (PBS), pH 7.4, containing 0.01 % sodium azide (Sigma) at 4°C until further processing for immunostaining. Once removed from the storing solution, the sections were rinsed in PBS and then incubated in 0.1 % Triton X-100 in PBS for 10 minutes at room temperature. To prevent nonspecific binding, the sections were blocked in a solution containing 5 % normal goat serum (NGS) (Vector Labs, Burlingame, CA), 0.1 % Triton X-100, 0.05 % Tween-20, and 1 % bovine serum albumin (BSA) for 30 minutes at room temperature. Subsequently, the sections were incubated for 72 hours at 4°C with specific primary antibodies for each experiment, including mouse anti-PKCδ (1:1000, BD Biosciences, 610397), rabbit anti-Phospho-c-Fos (1:2000, Cell Signaling Technology, 5348), rat anti-mCherry (1:500, Invitrogen, M11217) and rabbit anti-Phospho-p44/42 MAPK (Erk1/2) (1:200, Cell Signaling Technology, 9101L) These primary antibodies were diluted in a 1.5 % NGS blocking solution with 0.1 % Triton X-100, 0.05 % Tween-20, and 1 % BSA. Following the primary antibody incubation, the sections were rinsed in PBS and incubated with appropriate fluorescent secondary antibodies, such as Alexa Fluor 647-conjugated goat anti-mouse (1:100, Invitrogen, A21235), Alexa Fluor 647-conjugated goat anti-rabbit (1:250, Invitrogen, A21244), or goat anti-rat Cy3 (1:250, Invitrogen, A10522) in a 1.5 % NGS blocking solution with 0.1% Triton X-100, 0.05 % Tween-20, and 1 % BSA. This incubation step was performed for 2 hours at room temperature while protecting the sections from light. Finally, the sections were rinsed in PBS, mounted on positively charged glass slides, air-dried, and coverslips were placed using Fluoromount-G (Southern Biotech).

#### Image Acquisition and Quantification

For image acquisition, a Nikon A1R laser scanning confocal microscope was utilized, employing a 2X objective for low magnification, a 10X, 20X objective for general imaging, and a 40X oil-immersion objective for high magnification. All subsequent analyses were conducted using images obtained with the 10X and 20X objective, and the experimenter remained blinded to the experimental groups. Laser intensity, gain, and pinhole settings were maintained consistently across all experiments. Sequential image acquisition was conducted using multiple channels (GFP, RFP, and CY5), capturing Z stacks with a 0.9 mm interval. The acquired images were then processed using NIS Elements software, which automatically stitched them together and converted the image stacks into maximum-intensity z-projections. The quantitative analysis focused on the CeA region, specifically between the bregma coordinates of −0.70 and −2.06, with the aid of a mouse brain atlas (Paxinos and Franklin, 2001) to identify the anatomical boundaries. Manual cell quantification (automatic quantification for Fos experiment) was performed for each channel using NIS Elements, with one section analyzed per rostro-caudal level per mouse. Co-labeled cells were automatically identified by the software and further confirmed visually by the experimenter to ensure accuracy. Throughout the process, consistent methodologies were employed to maintain the reliability and validity of the findings.

### Statistical Analysis

All data are expressed as mean± SEM. Statistical analysis were performed were performed using two-way analysis of variance (ANOVA) followed by Tukey’s multiple comparison test, unpaired two-tailed t-test or linear regression, as appropriate using GraphPad Prism (v. 10.0). The n (number of animals), statistical test, and p values are presented in the figure legends. P < 0.05 was considered statistically significant.

## Results

### Fidelity and efficiency of Calcrl-Cre expression in the CeA

We sought to study the fidelity and efficiency of *Calcrl*-Cre expression in the CeA in both viral-infected and double-transgenic reporter mice. RNAScope fluorescence (*in situ* hybridization) and immunohistochemistry revealed that both groups of animals had similar fidelity of *Calcrl*-Cre expression. The mean percent of *Calcrl*-expressing CeA cells that also expressed the fluorescent maker was 83.5 ± 1.575 % in virus-infected mice and 84.8 ± 3.426 % in double transgenic reporter mice **(Fig. 1C)**. Next, we sought to determine how much of the endogenous *Calcrl* expression is targeted by either the virus or developmental tdTomato expression in double transgenic reporter mice by calculating efficiency. An unpaired t-test revealed the efficiency of *Calcrl*-Cre expression was significantly lower between virus-infected and double-transgenic reporter mice (p = 0.02). While virus-infected animals displayed a mean efficiency of 77.3 ± 11.9 %, transgenic reporter mice displayed a mean efficiency of 91.4 ± 1.71 %. This difference is attributed to the challenges of stereotaxic viral targeting **(Fig. 1C)**.

**Figure 1.**
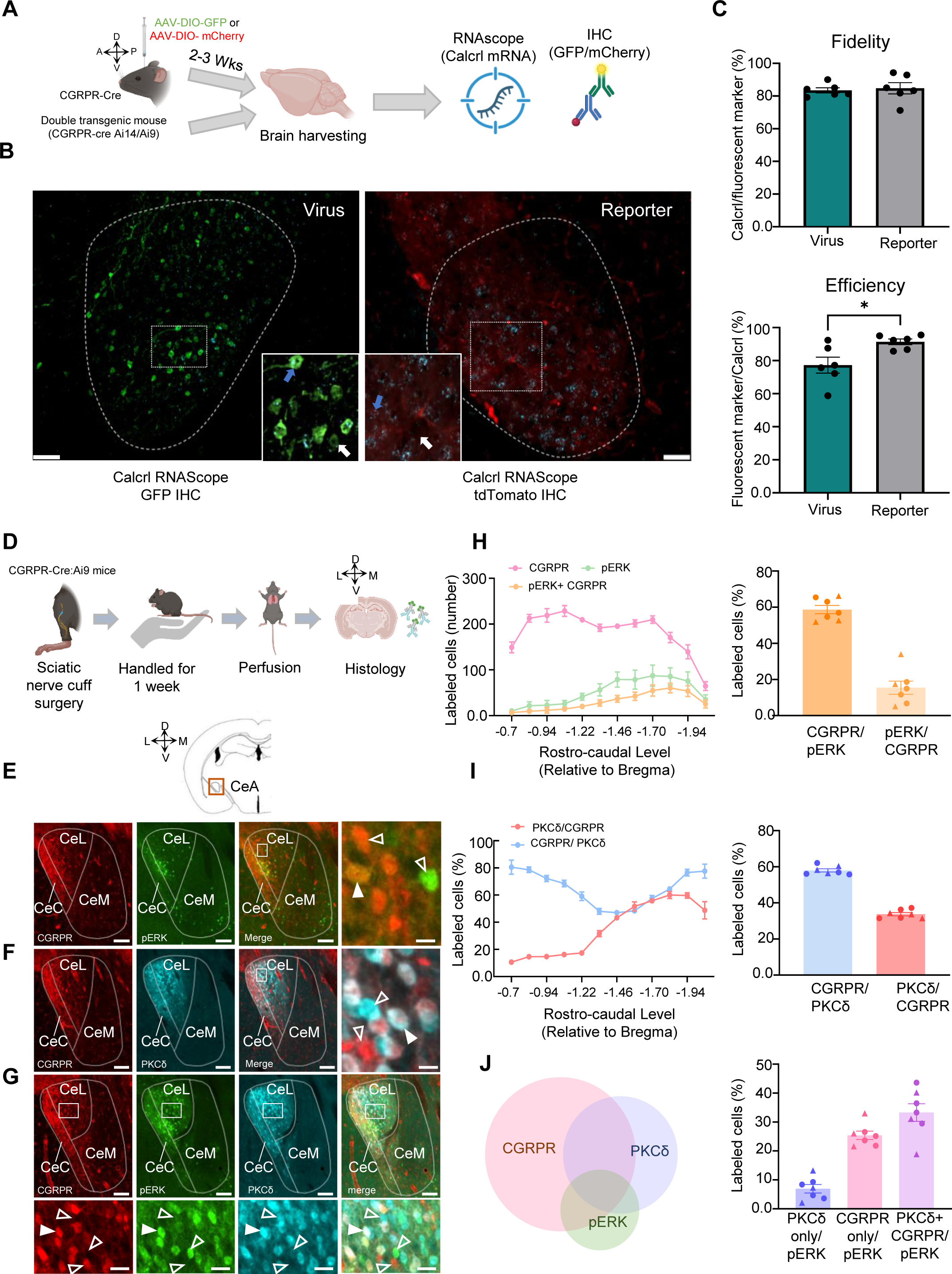
Nerve injury-induced pERK expression in CeA-CGRPR neurons. **(A)** Experimental timeline: Specificity of the Calcrl-Cre line is confirmed by colocalization of Calcrl mRNA with reporters from viral infection (AAV-DIO-GFP or AAV-DIO-mCherry) or transgenic lines (CalcrlCre:Ai14/Ai9). **(B)** Representative RNAScope/IHC images of the right CeA of a Calcrl-Cre animal injected with AAV-DIO-GFP (top) and a CalcrlCre:Ai9 double-transgenic animal (bottom) at 20x magnification and zoomed-in (inset) to show Calcrl-Cre positive (blue arrow, at least 3 puncta) and Calcrl-Cre negative (white arrow, < 3 puncta) cells within the CeA. Scale bar = 50 µm. **(C)** Quantification of fidelity (top; *Calcrl* positive mRNA divided by fluorescent marker multiplied by 100 %) and efficiency (bottom; fluorescent marker divided by Calcrl positive mRNA multiplied by 100%) of Calcrl-Cre expression in the CeA of viral-injected (teal bar) and double-transgenic reporter mice (gray bar). All data are presented as mean ± SEM. Each data point represents one CeA section. 2-3 sections were quantified in 3 different animals. An unpaired t-test revealed a significant difference in efficiency (*p < 0.05). **(D)** Experimental timeline for analysis of CeA-pERK in CGRPR reporter mice one week after sciatic nerve injury **(E-G)** Representative coronal CeA slices immunostained one week after nerve injury for pERK (green), CGRPR tdTomato (red), and PKCδ (cyan). **(E)** CGRPR and pERK; **(F)** CGRPR and PKCδ; **(G)** pERK, PKCδ and CGRPR. White open arrowheads indicate single labeled cells, and solid arrowheads indicate co-labeled cells. Scale bars represent 100 μm (low magnification) in the and 25 μm in the bottom high magnification images. **(H)** The rostral-caudal distribution of CGRPR and pERK labeled cells is shown on the left (n = 8 mice, 10 to 12 slices per mouse), and the mean ± SEM percentage of cells co-labeled for pERK and CGRPR is shown on the right (n = 7 mice, 11 slices per mouse). **(I)** The left panel shows the rostral-caudal distribution of CGRPR and PKCδ labeled cells (n = 8 mice, 10 to 12 slices per mouse), and the mean ± SEM percentage of cells co-labeled for pERK and CGRPR is shown on the right (n = 7 mice, 11 slices per mouse). **(J)** Pie chart showing distribution and overlap of the CGRPR, PKCδ, and pERK positive cell populations in the CeA one week after nerve injury. The mean ± SEM percentage of cells co-labeled for pERK, CGRPR, and PKCδ positive cells are shown on the right (n = 7 mice, 11 slices per mouse). The data points represented by triangles on the graph correspond to female mice.

### Nerve injury-induced pERK expression in CeA-CGRPR neurons

Phosphorylated extracellular signal-regulated kinase 1/2 (pERK) plays a significant role in the development of persistent pain and serves as a marker for central sensitization. Previous research has shown that the activation pERK by the upstream kinase in pain pathways, such as the dorsal horn and CeA neurons (Dai et al., 2002; Zhuang et al., 2005; Carrasquillo and Gereau 2007; Wilson et al., 2019). In our study, we conducted immunostaining analysis and observed a significant increase in the number of pERK-positive cells co-expressing the CGRPR protein in the CeA one week after cuff implantation in the sciatic nerve **(Fig. 1E)**. These CGRPR-positive neurons were distributed throughout different rostro-caudal levels of the CeA, with a particular concentration in the medial and posterior areas **(Fig. 1H)**. Further analysis revealed that 58.6 % of the total pERK-positive cells co-localized with CGRPR-positive cells, whereas only 15.8 % of the total CGRPR-positive cells showed co-labeling with pERK-positive cells **(Fig. 1H)**. A study from our lab (Wilson et al., 2019) demonstrated that ERK activation induced by nerve injury is preferential localized to neurons expressing protein kinase C delta (PKCδ) in the CeA. To further investigate this phenomenon, we performed immunolabeling of PKCδ cells in the same CGRPR-Ai9 mice **(Fig. 1F-G)** and conducted a quantitative analysis.

Notably, we observed a significant overlap between CGRPR and PKCδ positive neurons in the medial and posterior regions of the CeA. Further analysis indicated that 58 % of PKCδ neurons expressed the CGRPR protein, while 33 % of CGRPR neurons expressed the PKCδ protein **(Fig. 1I)**. Subsequently, we conducted a comprehensive analysis of pERK labeling within the CGRPR and PKCδ neuronal populations in the CeA. Our findings revealed that 6.93 % of pERK was exclusively detected in cells expressing PKCδ, while 25.39 % of pERK was solely observed in cells expressing CGRPR. Furthermore, we observed that 33.27 % of pERK was present in cells that co-expressed both CGRPR and PKCδ proteins **(Fig. 1J)**. Further, characterization revealed no differences in the number of CGRPR or PKCδ neuronal populations across rostro-caudal level, nor in their colabeling, between males and females across the left and right hemisphere **(Fig. S1)**. These results suggest a potential functional interaction between CGRPR and PKCδ signaling in the CeA, which may be related to pain processing and central sensitization.

### Intrinsic excitability of CGRPR expressing neurons in a neuropathic mice model of pain is dependent on rostro-caudal level

Previous studies have indicated the abundant expression of CGRP in PBN neurons projecting to the CeA (D’Hanis, Linke et al., 2007; Carter., et al., 2013). Stimulation of these PBN axonal fibers results in CeA neuron depolarization, suggesting induced synaptic plasticity (Han, Li et al., 2005; Han, Adwanikar et al., 2010). Genetic inhibition of CGRPR neurons has been found to attenuate fear response (Han et al., 2015). However, there is limited research on the intrinsic excitability of CGRPR-expressing CeA neurons in acute pain models. Hence, we conducted patch clamp electrophysiology to evaluate the intrinsic excitability of these neurons in a neuropathic pain model **(Fig. 2A)**.

**Figure 2.**
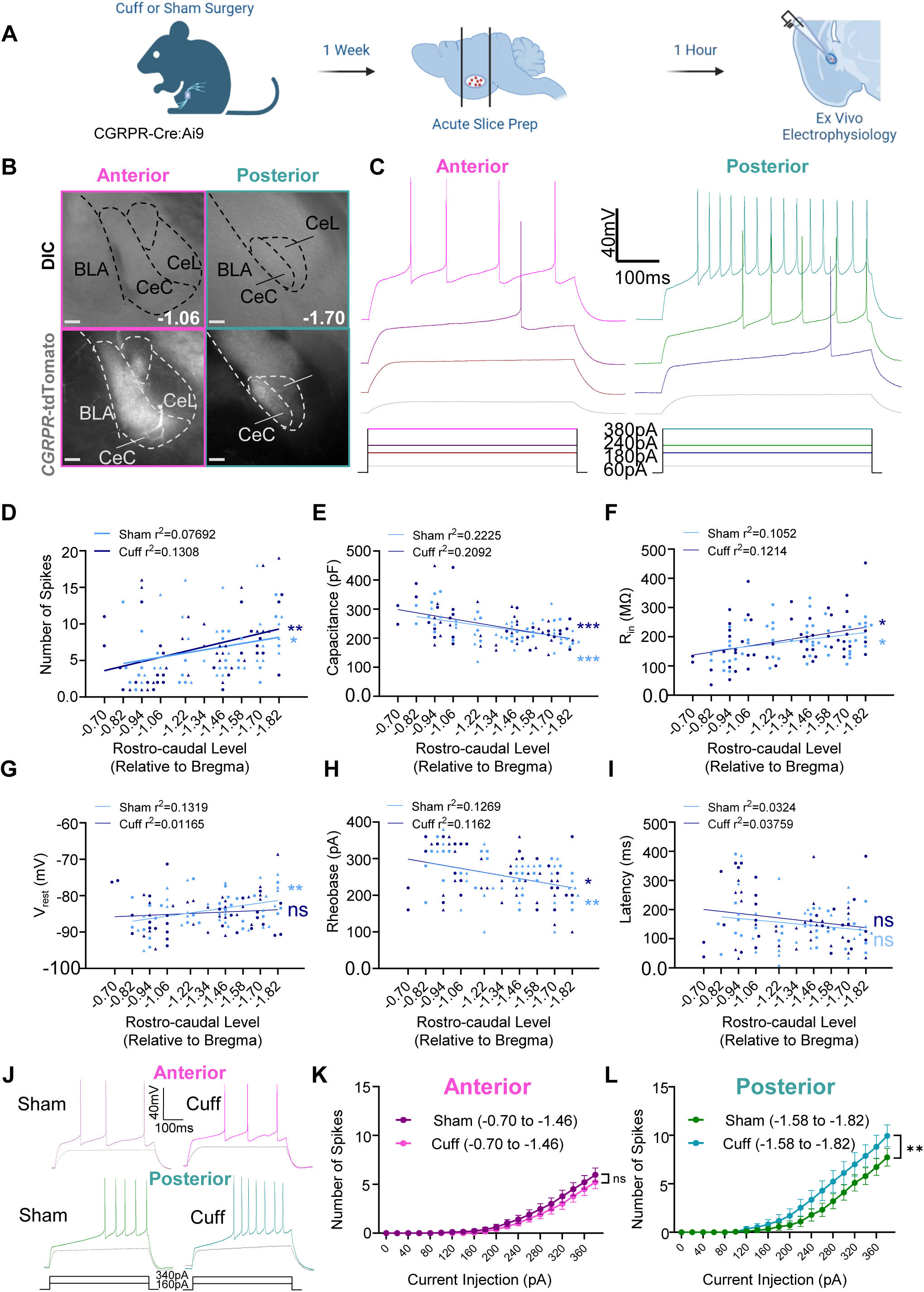
Intrinsic excitability of CGRPR expressing neurons in a neuropathic model of pain is dependent on rostro-caudal level. **(A)** Diagram of experimental procedure and timeline. *CGRPR-cre*:Ai9 mice were handled for at least 5 days leading up to acute slice preparation. Acute slice preparation occurred following transcardial perfusion with a choline chloride-based cutting solution. **(B)** Examples of acute slices comparing anatomical landmarks at relevant rostro-caudal levels. (Left) Representative images of an anterior (-1.06 mm relative to bregma) rostro-caudal level. Differential interference contrast images (top) show fiber tracts and landmarks used to identify the amygdala, while fluorescent images (bottom) show corresponding tdTomato-expressing CGRPR neurons. (Right) Differential interference contrast image (left) and fluorescent image (right) of a posterior (-1.70 mm relative to bregma) rostro-caudal level. **(C)** Representative voltage traces comparing one anterior (-1.06 mm relative to bregma) and one posterior (-1.82 mm relative to bregma) CeA-CGRPR cells. Traces correspond to increasing current injection steps of 60, 180 (rheobase for the posterior neuron), 240 (rheobase for the anterior neuron), and 380 pA. **(D)** Scatterplot comparing the number of action potentials elicited by a 500 ms current injection of 380 pA (the greatest amount of current injected which did not fail) with rostro-caudal level. Both sham (p=0.0465, r^2^=0.07692) and cuff (p=0.0078, r^2^=0.1308) treatments show a correlation between the rostro-caudal level and the number of action potentials after a simple linear regression. Data recorded from female animals is indicated by triangles while data from male animals is indicated by circles. Sham male n = 25, Sham female n = 27. Cuff male n = 27, Cuff female n = 26. **(E)** Scatter plot comparing membrane capacitance with rostro-caudal level. Sham (p=0.0004, r2=0.2225P) and Cuff (p=0.0006, r2=0.2092) significantly correlate with rostro-caudal level after a simple linear regression. Data recorded from female animals is indicated by triangles while data from male animals is indicated by circles. Sham male n = 25, Sham female n = 27. Cuff male n = 27, Cuff female n = 26. **(F)** Scatter plot comparing input resistance with rostro-caudal level. Both Sham (p=0.0190, r^2^=0.1052) and Cuff (p=0.0106, r^2^=0.1214) show a significant correlation between input resistance and rostro-caudal level, Data recorded from female animals is indicated by triangles while data from male animals is indicated by circles. Sham male n = 25, Sham female n = 27. Cuff male n = 27, Cuff female n = 26. **(G)** Scatter plot comparing resting membrane potential with rostro-caudal level. Sham cells show a correlation between resting membrane potential and rostro-caudal level (p=0.0081, r^2^=0.1319), but cuff cells do not (p=0.4416, r^2^=0.01165). Data recorded from female animals is indicated by triangles while data from male animals is indicated by circles. Sham male n = 25, Sham female n = 27. Cuff male n = 27, Cuff female n = 26. **(H)** Scatter plot comparing rheobase with rostro-caudal level. Both Sham (p=0.0096, r^2^=0.1269) and Cuff conditions (p=0.0125, r^2^=0.1162) show a correlation between rheobase and rostro-caudal level. Data recorded from female animals is indicated by triangles while data from male animals is indicated by circles. Sham male n = 25, Sham female n = 27. Cuff male n = 27, Cuff female n = 26. **(I)** Scatter plot comparing latency to the first spike at 380 pA with rostro-caudal level. Cells recorded from both sham (p=0.2016, r^2^=0.03241) and cuff (p=0.1642, r^2^=0.03759) mice show a correlation between latency and rostro-caudal level. Data recorded from female animals is indicated by triangles while data from male animals is indicated by circles. Sham male n = 25, Sham female n = 27. Cuff male n = 27, Cuff female n= 26. **(J)** Representative voltage traces of CeA-CGRPR cells in either cuff or sham mice. (Top) Comparison of sample traces collected from anterior rostro-caudal levels between cuff and sham animals. Firing is shown at two current injection steps for each: 220 pA (color), and 160 pA (gray). (Bottom) Comparison of sample traces collected from posterior rostro-caudal levels between cuff and sham animals. Firing is shown at two current injection steps for each: 220 pA (color), and 160 pA (gray). **(K)** Plot of action potentials fired at each current step by anterior neurons (-0.70 to -1.46 mm relative to bregma) in cuff and sham conditions. The number of action potentials evoked is plotted as a function of the respective amplitudes of the current injected. All error bars represent the standard error of the mean (SEM). This graph includes combined data from male and female mice. Results of two-way ANOVA: Effect of current injection, p < 0.0001; effect of treatment, p = 0.3992 ns; interaction, p = 0.9547 ns (n = 34 neurons for sham and 36 for cuff). **(L)** Plot of action potentials fired at each current step by posterior neurons (-1.58 to -1.82 mm relative to bregma) in cuff and sham conditions. The number of action potentials evoked is plotted as a function of the respective amplitudes of the current injected. All error bars represent the standard error of the mean (SEM). This graph includes combined data from male and female mice. Results of two-way ANOVA: Effect of current injection, p < 0.0001; effect of treatment, p = 0.1613 ns; interaction, p = 0.0055 (n = 19 neurons for sham and 17 for cuff). The data points represented by triangles on the graph correspond to female mice.

Due to the functional variability of CGRPR neurons in the CeA across rostro-caudal levels (Bowen et al., 2023), we collected data from CGRPR-expressing CeA neurons at multiple rostro-caudal levels. Plotting excitability in response to a 500 ms, 380 pA current injection against the rostro-caudal level revealed a positive correlation in both cuff and sham conditions, with higher excitability observed in posterior levels compared to anterior levels **(Fig. 2D)**. Analysis of capacitance **(Fig. 2E)** reveals a significant relationship between the capacitance of both cuff and sham neurons and rostro-caudal level, with more posterior cells showing lower capacitance. Comparisons of capacitance between cuff and sham treatments in cells classified as anterior and posterior show no significant difference **(Table 1)**. As shown previously in other genetic subpopulations of neurons within the CeA (Adke, Khan, et al. 2021), a strong negative correlation between capacitance and excitability suggests that smaller neurons tend to be more excitable. These results suggest that capacitance plays a role in the excitability gradient across rostro-caudal levels but is less likely to contribute to the increased excitability observed in posterior cuff neurons. We also examined subthreshold membrane properties. Analysis of input resistance **(Fig. 2F)** shows a correlation with rostro-caudal level in both cuff and sham treatments, with higher Rin observed in cells at more posterior RC levels. The Rin of anterior and posterior neurons does not show a significant difference between cuff and sham treatments **(Table 1)**. This suggests that conductance may contribute to the overall increased excitability of neurons at posterior rostro-caudal levels.

**Table 1.**
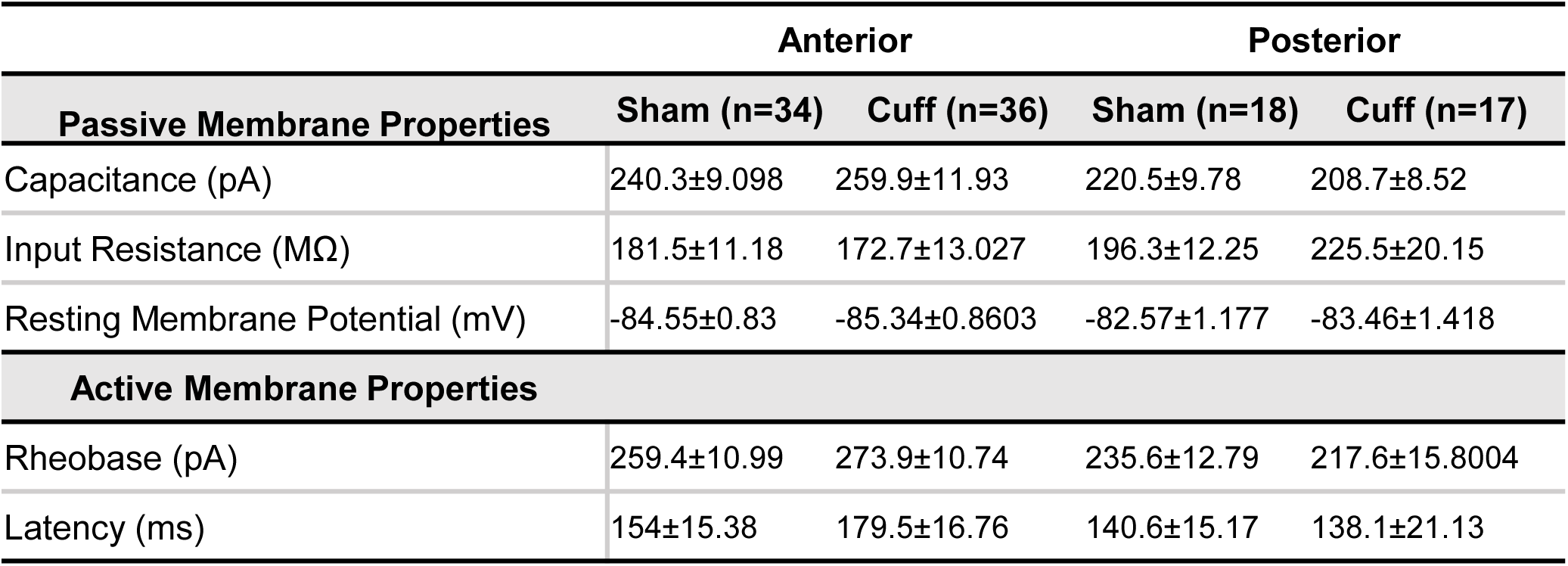
Passive and active membrane properties of CGRPR-CeA neurons from anterior or posterior rostro-caudal levels in a mouse model of neuropathic pain. All data is presented as means ± SEM. Unpaired t-test comparing anterior or posterior cuff and sham conditions.

Additionally, analysis of Vrest **(Fig. 2G)** shows a correlation with RC level in the sham condition only, with more depolarized resting membrane potential observed at posterior RC levels, while the Vrest of neurons in the cuff condition does not correlate with RC level. When examining the differences between cuff and sham neurons from anterior and posterior RC levels, we see no differences in Vrest **(Table 1)**. This suggests that more depolarized Vrest is associated with the greater excitability observed in neurons found in posterior rostro-caudal levels, however, it is unlikely that this contributes to the increased excitability of posterior cuff neurons.

We also chose to plot suprathreshold properties such as rheobase and latency to fire as a function of rostro-caudal level, and we see a correlation between rheobase and rostro-caudal level in both cuff and sham conditions **(Fig. 2H)**. Latency does not show a correlation with rostro-caudal level in either condition **(Fig. 2I)**. Rheobase and latency do not appear to be significantly different between cuff and sham conditions in either anterior or posterior neurons **(Table 1)**.

When plotting the excitability of CeA CGRPR neurons against increasing current injection amplitudes (0-380 pA), we observed increased excitability in response to higher current amplitudes **(Fig. 2K-L)**. Further analysis showed no change in excitability between cuff and sham treatments in cells classified as anterior **(Fig. 2K)**. However, a two-way analysis of variance (ANOVA) revealed a significant (p < 0.0001) increase in excitability in the cuff condition at posterior rostro-caudal levels **(Fig. 2L)**. These findings establish a relationship between post-synaptic properties and rostro-caudal level in CGRPR-expressing neurons within the CeA.

### Inhibition of CeA-CGRPR neurons reverses cuff-induced hypersensitivity

The results obtained from pERK activation and *ex vivo* electrophysiology strongly support the notion that the sensitization of CeA-CGRPR neurons plays a vital role in pain modulation. To establish a causal relationship between the heightened activity of CeA-CGRPR neurons and pain-related behavior, we employed a chemogenetic strategy in a sciatic nerve cuff/sham mice model (Benbouzid et al. 2008). We performed stereotaxic injections of cre-dependent inhibitory DREADD hM4Di or control mCherry virus into the CeA of CGRPR-Cre mice. As demonstrated in **Figures 3B-C**, the unilateral stereotaxic administration of hM4Di into the CeA yielded strong and localized transduction of hM4Di-mCherry within CeA-CGRPR cells. Additionally, transduced cell mapping validated the precise localization of the injection sites within the CeA region **(Fig. 3D)**. To verify the efficacy of chemogenetic manipulation, we conducted immunohistochemical analysis targeting Fos, a well-established marker for neuronal activation **(Fig. 3E)**. As depicted in **Figure 3F**, administration of CNO via intraperitoneal injection (i.p) resulted in a significant reduction in Fos expression within the CeA of CGRPR-cre mice transduced with hM4Di, as compared to control group of mice expressing mCherry. This observation is further supported by quantitative analysis of co-localized Fos and mCherry cells within the CeA, reaffirming a significantly (p < 0.01) lower Fos expression in hM4Di-transduced cells among mice treated with CNO, compared with mCherry controls **(Fig. 3G)**.

**Figure 3.**
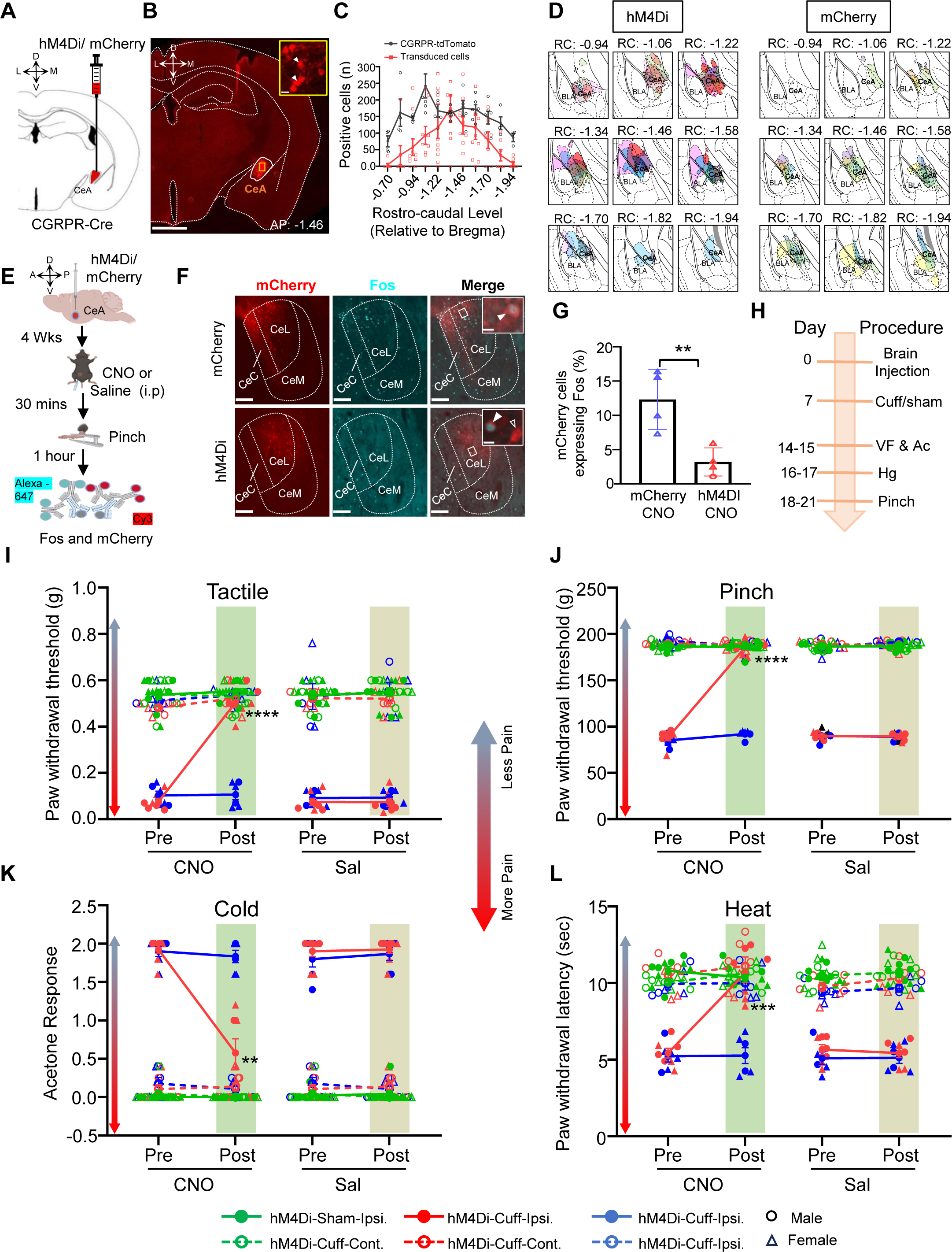
Inhibition of CeA-CGRPR neurons reverses cuff-induced hypersensitivity. **(A)** Schematic diagram for unilateral stereotaxic injection of hM4Di or mCherry into the right CeA of CGRPR-cre mice. **(B)** A representative image of a coronal mouse brain slices from a CGRPR-cre mouse infected with hM4Di into the CeA. Scale bar: 1000 µm. The transduced cells in area delineated by the yellow box in CeA are shown at higher magnification in inset panel. White arrowheads indicate cells positive for hM4Di. Scale bar: 20 µm. **(C)** Mean ± SEM number of hM4Di and mCherry transduced cells and CGRPR-tdTomato labeled cells in the CeA as a function of rostral-caudal level relative to bregma. N = 10 (4 male and 6 female mice) for transduced neurons and n = 6 (3 male and 3 female mice) for CGRPR-tdTomato neurons. **(D)** Rostral-Caudal map of CeA neurons transduced with hM4Di or mCherry in CGRPR-Cre mice used in study. Each color represents the viral expression spread for an individual animal. **(E)** Experimental timeline for validating hM4Di-mediated inhibition of pinch-induced Fos expression in CeA-CGRPR neurons. **(F)** Representative images of coronal brain slices containing the CeA of CGRPR-cre mice infected with mCherry (top panel) or hM4Di (lower panel) into the CeA and i.p. treated with CNO. hM4Di or mCherry transduced cells are shown in red, and immunostaining for Fos is shown in cyan. The merged images are shown in the right panels. Inset white boxes delineate the areas magnified in the rightmost panel. White solid arrowheads point to cells positive for both transduced cells and Fos, while open arrowheads point to cells that are positive for hM4Di or mCherry only. Scale bars: 100 μm (low magnification) and 10 μm for inset magnified images. **(G)** Mean ± SEM numbers of Fos and mCherry transduced co-labeled cells per condition. Unpaired two-tailed t test (n = 4 per group; 3 females and 1 male per group), p = 0.0095. Data from female animals is indicated by triangles. **(H)** Timeline for behavioral experiments. **(I-L)** Battery of pain behavior test. Results are shown as mean ± SEM for hypersensitivity in ipsilateral and contralateral hindpaw before and 1-hour after CNO (or saline) i.p. injection in CGRPR-cre mice stereotaxically infected with hM4Di or mCherry into the CeA. Repeated-measure two-way ANOVA followed by Tukey’s multiple comparison test was performed for analysis of all behavioral assays. **(I)** Von-Frey test to measure tactile hypersensitivity. P < 0.0001 for pre CNO vs post CNO in Cuff-Gi Ipsilateral. **(J)** Randall-Sellitto test to measure pinch hypersensitivity. P < 0.0001 for pre CNO vs post CNO in Cuff-Gi Ipsilateral. **(K)** Acetone test to measure cold hypersensitivity. P = 0.0034 for pre CNO vs post CNO in Cuff-Gi Ipsilateral. **(L)** Hargreaves test to measure heat hypersensitivity. P = 0.0004 for pre CNO vs post CNO in Cuff-Gi Ipsilateral. N = 10 (5 females and 5 males) for hM4Di-sham; n = 7 (3 females and 4 males) for hM4Di-cuff; and n = 6 (4 females and 2 males) for mCherry-cuff animals. Individual mice are represented by scatter points and female mice are labeled in triangles.

Next, we examined the impact of targeted chemogenetic inhibition of CeA-CGRPR neurons on pain-related behavior to tactile, pinch, cold, and heat stimuli applied to the hindpaws **(Fig. 3H)**. We assessed behavioral responses both before and after i.p administration of either saline or CNO (10 mg/kg) in both the cuff and sham groups of mice. In the left hindpaw ipsilateral to cuff implantation, the selective inhibition of right CGRPR-positive CeA neurons using CNO led to a significant reversal of cuff-induced tactile hypersensitivity (p < 0.0001) and pinch hypersensitivity (p < 0.0001). However, no measurable effects were observed in saline-injected or CNO-injected mCherry-control mice **(Fig. 3I-J)**. **Figure 3K-L** shows that the reversal of cuff-induced hypersensitivity was not limited to a particular modality, as the chemogenetic inhibition of CGRPR-positive neurons in the CeA also reduced hypersensitivity responses to cold (p < 0.01) and heat (p < 0.001) stimulation of the hindpaws. Notably, the impact of inhibiting CGRPR-positive CeA neurons was restricted to nerve injury, as stimuli-induced response in sham-treated mice or the hindpaw contralateral to the sciatic nerve cuff remained comparable among all experimental groups. Taken together, these results demonstrate that inhibiting CGRPR-positive neurons in the CeA effectively reverses cuff-induced pain hypersensitivity.

### Activation of CeA-CGRPR neurons is sufficient to induce pain-related hypersensitivity without injury

Our subsequent experiments aimed to determine whether the activation of CeA-CGRPR neurons alone could induce pain-related hypersensitivity in the absence of injury. To achieve this, we injected the excitatory DREADD hM3Dq into the right CeA of CGRPR-cre mice **(Fig. 4A-B)** and subjected them to a battery of pain behavioral tests before and after an i.p. injection of CNO (5 mg/kg) or saline, in both the cuff and sham groups. The unilateral stereotaxic administration of hM3Dq into the CeA yielded strong and localized transduction of hM3Dq-mCherry within CeA-CGRPR cells across rostro-caudal level **(Fig. 4C)**. Histological analysis confirmed the accurate targeting of hM3Dq transduction within the CeA, as observed by the restricted transduction of hM3Dq in the CeA region **(Fig. 4D)**. **Figure 4E-G** shows that i.p. administration of CNO robustly induced Fos expression in the CeA of CGRPR-cre mice infected with hM3Dq, compared to saline-injected control mice **(Fig. 4F)**. Quantification of co-expressing Fos and mCherry cells in the CeA confirmed significantly higher Fos expression (p < 0.01) in hM3Dq-transduced cells with CNO treatment **(Fig. 4G)**.

**Figure 4.**
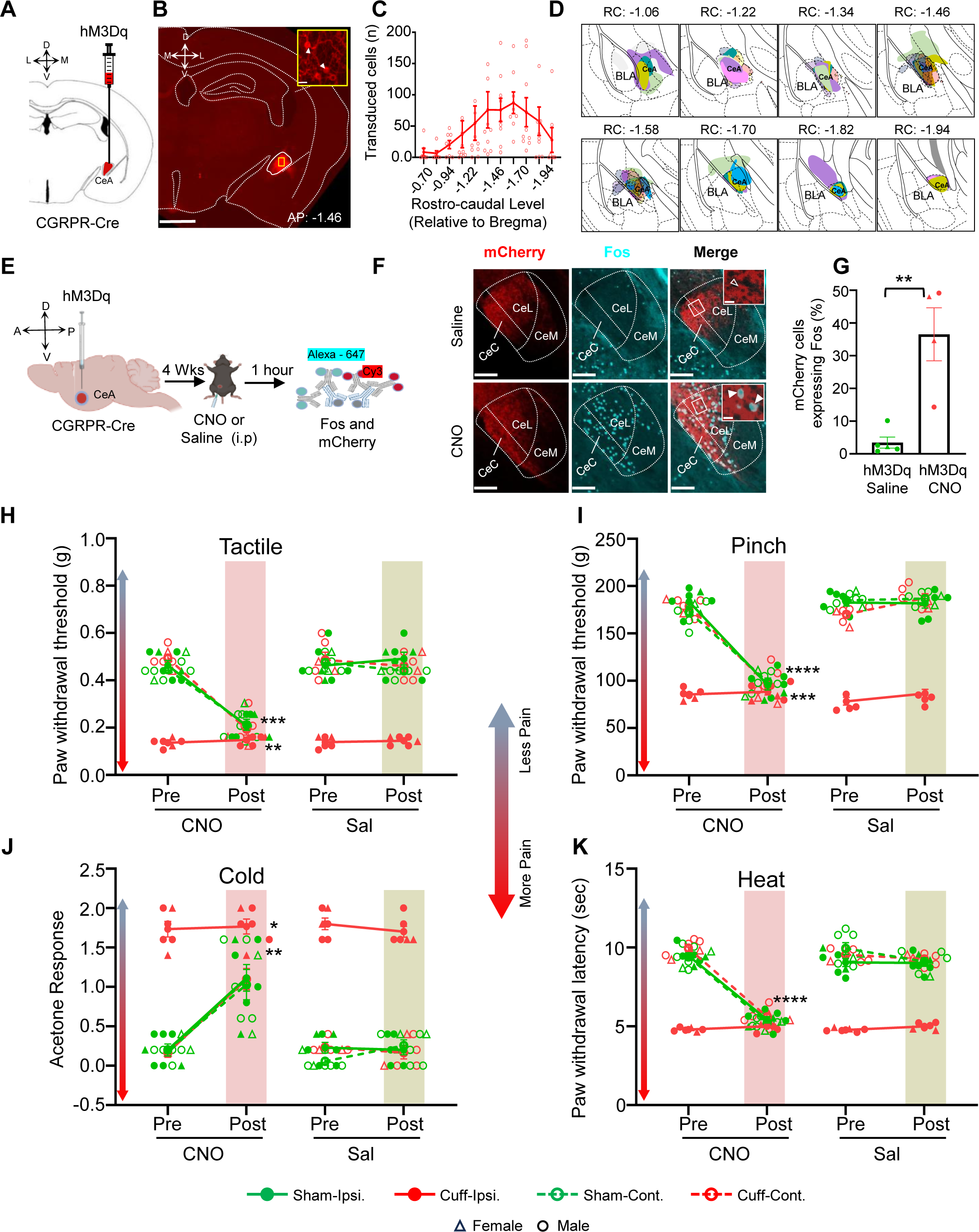
Activation of CeA-CGRPR neurons induces pain-related hypersensitivity without injury. **(A)** The schematic diagram for unilateral stereotaxic injection of hM3Dq into the CeA of CGRPR-cre mice. **(B)** A representative image of a coronal mouse brain slices from a CGRPR-cre mouse infected with hM3Dq into the CeA. Scale bar: 1000 µm. The area delineated by the yellow box is shown at higher magnification in inset panel. The transduced cells in area delineated by the yellow box in CeA are shown at higher magnification in inset panel. White arrowheads indicate cells positive for hM3Dq. Scale bar: 20 µm. **(C)** Mean ± SEM number of hM3Dq-transduced cells in the CeA as a function of rostral-caudal level relative to bregma (n = 10 mice for hM3Dq-transduced neurons). **(D)** Drawings depicting the rostral-caudal distribution of hM3Dq injection sites in mice used for experiments. Individual mice are represented by different colors. **(E)** Experimental timeline for validating hM3Dq-mediated activation of Fos expression in CeA-CGRPR neurons. **(F)** Representative images of coronal brain slices containing the CeA of CGRPR-cre mice infected with hM3Dq into the CeA and i.p. treated with saline (top panel) or CNO (bottom panel). hM3Dq transduced cells are shown in red, and immunostaining for Fos is shown in cyan. The merged images are shown in the right panels. White boxes delineate the areas magnified in the inset panel. White open arrowheads point to cells that are positive for hM3Dq only, while white solid arrowheads point to cells positive for both transduced cells and Fos. Scale bars: 100 μm (low magnification) and 10 μm for inset magnified images. **(G)** Mean ± SEM numbers of Fos and mCherry transduced co-labeled cells per condition. Unpaired two-tailed t test (Gq-saline: n = 5, 1 female and 4 males; Gq-CNO: n = 4, 2 female and 2 male), p = 0.0028. **(H-K)** Battery of pain behavior test. Results are shown as mean ± SEM for hypersensitivity in ipsilateral and contralateral hindpaw before and 1-hour after CNO (or saline) i.p. injection in CGRPR-cre mice stereotaxically infected with hM3Dq into the CeA. Repeated-measure two-way ANOVA followed by Tukey’s multiple comparison test was performed for analysis of all behavioral assays. **(H)** Von-Frey test to measure tactile hypersensitivity. Post hoc comparisons revealed significant differences between pre-CNO and post-CNO in Sham-Gq ipsilateral (p = 0.0003), Sham-Gq contralateral (p = 0.0015), and Cuff-Gq contralateral (p = 0.0009). **(I)** Randall-Sellitto test to measure pressure hypersensitivity. Post hoc comparisons revealed significant differences between pre-CNO and post-CNO in Sham-Gq ipsilateral (p < 0.0001), Sham-Gq contralateral (p < 0.0001), and Cuff-Gq contralateral (p = 0.0006). **(J)** Acetone test to measure cold hypersensitivity. Post hoc comparisons revealed significant differences between pre-CNO and post-CNO in Sham-Gq ipsilateral (p = 0.0182), Sham-Gq contralateral (p = 0.0297), and Cuff-Gq contralateral (p = 0.0080). **(K)** Hargreaves test to measure heat hypersensitivity. Post hoc comparisons revealed significant differences between pre-CNO and post-CNO in Sham-Gq ipsilateral (p < 0.0001), Sham-Gq contralateral (p < 0.0001), and Cuff-Gq contralateral (p < 0.0001). n = 7 (2 females and 5 males) mice for sham-hM3Dq, and n = 6 (2 females and 4 males) mice for cuff-hM3Dq. Individual mice are represented by scatter points and female mice are identified in triangles.

At the behavioral level, chemogenetic activation of CeA-CGRPR neurons induced hypersensitivity to tactile, pinch, cold and heat stimuli even in the absence of nerve injury. Significant pre- vs post-CNO differences were observed in sham animals for tactile (left hindpaw ipsilateral to sham injury: p = 0.0003; right hindpaw contralateral to sham injury: p = 0.0015), pinch (left hindpaw ipsilateral to sham injury: p < 0.0001; right hindpaw contralateral to sham injury: p < 0.0001), cold (left hindpaw ipsilateral to sham injury: p = 0.00182; right hindpaw contralateral to sham injury: p = 0.00297), and heat (left hindpaw ipsilateral to sham injury and right hindpaw contralateral to sham injury: p < 0.0001) responses. in cuff animals, CNO increase hypersensitivity in the right hindpaw contralateral to cuff injury across tactile (p = 0.0009), pinch (p = 0.0006), cold (p = 0.0080), ad heat (p < 0.0001) tests, whereas hypersensitivity in the injured paw (left hindpaw ipsilateral to injury) was not affected, possibly due to the mice already reaching maximum levels of hyperalgesia **(Fig. 4H-K)**. Collectively, these findings highlight the essential role of CeA-CGRPR neuron activity in peripheral hypersensitivity regardless of the sensory modality and demonstrate that their activation in the absence of injury alone is sufficient to induce hypersensitivity.

### Chemogenetic inhibition of CeA-CGRPR neurons reduces formalin-induced spontaneous pain responses only in female mice

We investigated the role of CeA-CGRPR neurons in spontaneous nociceptive responses using the well-established formalin model of inflammatory pain (Abbott et al., 1995) in the context of DREADD inhibition of right CeA CGRPR expressing neurons. Nociceptive behaviors were evaluated following the CNO systemic injection and subcutaneous injection of 10 µL of 3 % formalin into the plantar surface of either the left or right hind paw. Our findings revealed sex-specific differences in formalin behavior following the inhibition of CeA-CGRPR neurons. The chemogenetic inhibition of CeA-CGRPR neurons in female mice led to a significant reduction in all cumulative nociceptive behaviors (p < 0.001) compared to control mice **(Fig. 5B)**. However, no measurable effects were observed in CNO-injected male mice **(Fig. 5C)**. Further analysis revealed that mice did not show any significant differences in the first phase of formalin response in either female **(Fig. 5D)** or male mice **(Fig. 5E)**. However, female mice treated with CNO exhibited a significant (p < 0.01) decrease in the second phase of formalin-induced nociceptive behaviors compared to control mice **(Fig. 5F)**. Conversely, male mice didn’t show any significant difference in the second phase of formalin-induced nociceptive behavior **(Fig. 5G)**. Altogether, these results indicate sex-specific differences in formalin-induced inflammatory spontaneous pain response.

**Figure 5.**
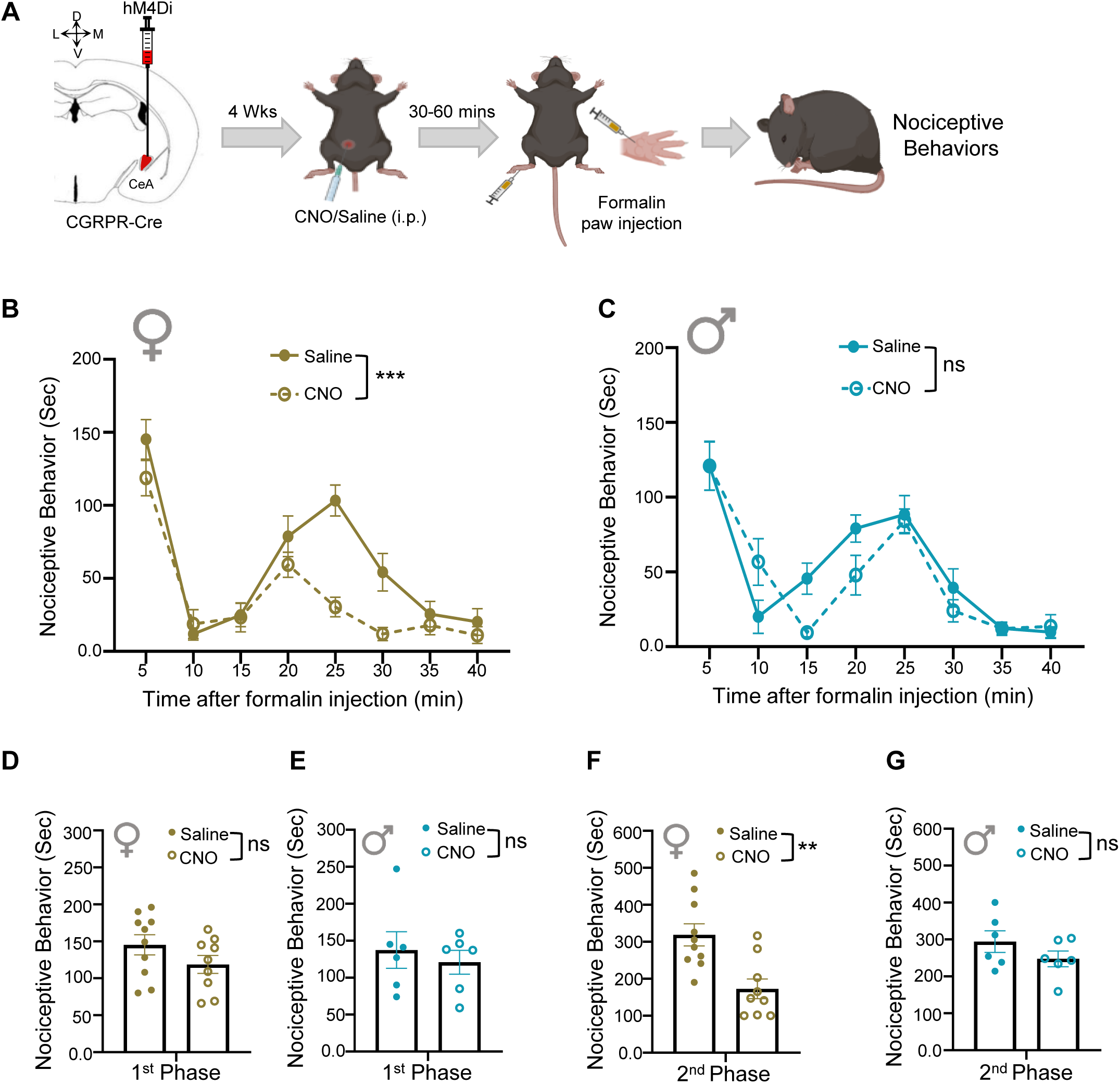
Inhibition of CGRPR expressing neurons in CeA reduces formalin-induced spontaneous pain behavior only in female mice. **(A)** Experimental Timeline. After 4 weeks of viral infections, mice were injected (i.p) with either CNO or saline. After 30-60 minutes, the left or right hind paw was injected with 3 % formalin, and nociceptive behaviors were measured. **(B-C)** Time spent in nociceptive behaviors as a function of time (per 5 minutes interval) after formalin injection. **(B)** Female CGRPR cre mice. Repeated measures two-way ANOVA analysis revealed significant effects of i.p. treatment (F(1,17) = 16.01, p = 0.0009), time (F(3.512,59.70) = 36.29, p < 0.0001), and treatment × time interaction (F(3.512,59.70) = 3.688, p = 0.0124). **(C)** Male CGRPR cre mice. Repeated measures two-way ANOVA analysis revealed no significant effects of i.p. treatment (F(1,10) =1.344, p = 0.2732), a significant effect of time (F(3.772,37.72) = 25.71, p < 0.0001), and no significant effect treatment × time interaction (F(3.772,37.72) = 2.234, p = 0.0870). **(D-E)** Time spent in nociceptive behaviors Phase 1, which encompasses 0-5 minutes post-formalin injection in **(D)** Female and **(E)** Male mice. **(F-G)** Phase 2, which encompasses 5-40 minutes post-formalin injection. **(F)** Female and **(G)** Male mice. An unpaired two-tailed t-test was performed for analysis. For female mice phase 1: t = 1.458, df = 16.95; p = 0.1630; and phase 2: t = 3.644, df =16.92; p = 0.0020; for male mice 1st phase: t = 0.5574, df = 8.595; p = 0.5915; and phase 2: t = 1.289, df = 9.172; p = 0.2291. Female- n = 10 for saline and n = 9 animals for CNO groups; Male- n = 6 for each group. Individual mice are represented by scatter points. All data are presented as means ± SEM.

## Discussion

The CeA contains heterogenous GABAergic neurons with distinct protein expressions and is recognized for its association with nociceptive behavior. Recently, considerable attention has been directed towards CGRPR-expressing neurons in the CeA due to their involvement in CGRP-mediated pain modulation (Xu et al., 2003; Han et al., 2005, 2010; Allen et al., 2023; Trail et al., 2026). These neurons are targeted by CGRP neuropeptide, signifying their importance in pain-related processes within the CeA. However, these neurons’ specific functional properties and contributions to pain processing are unclear. Our findings demonstrate that neurons expressing CGRPR in the CeA exhibit a notable increase in phosphorylated ERK in response to nerve injury. Additionally, a subset of these neurons co-label with PKC-δ positive neurons in the CeA. Next, we evaluate the fidelity and efficiency of Calcrl-Cre expression in the CeA using two approaches: viral infection and double-transgenic reporter mice. The results demonstrate that both approaches reliably target Calcrl-expressing cells. Using *ex vivo* electrophysiological recordings, we demonstrated that CeA-CGRPR neurons showed changes in intrinsic excitability at the RC level following sciatic nerve cuff implantation. We further showed that chemogenetic inhibition of CeA-CGRPR neurons effectively reverses nerve injury-induced hypersensitivity. Conversely, activating these neurons induces bilateral hypersensitivity even in the absence of nerve injury. Our data shows the important role played by CeA-CGRPR positive neurons in pain processing and their contribution to understanding the neural circuits involved in persistent pain conditions.

### Nerve injury-induced recruitment and interplay of CGRPR, PKCδ, and pERK-expressing neurons in the central nucleus of the amygdala

The activation of pERK is known to play a crucial role in various cellular processes, including pain modulation (Carrasquillo and Gereau 2007; Wilson et al., 2019). In our study, we observed a significant increase in pERK-positive cells in the CeA one week after sciatic nerve cuff implantation, consistent with previous research demonstrating pERK upregulation in other pain pathways (Zhuang et al., 2005). The co-localization of pERK with CGRPR-positive cells in the CeA shows the involvement of CGRPR-expressing neurons in the activation of the ERK pathway following nerve injury. Additionally, we investigated the potential interplay between CGRPR and PKCδ signaling in the CeA as previous evidence has linked PKCδ to ERK activation in response to nerve injury (Wilson et al., 2019). In line with other studies, we observed a significant overlap between CGRPR and PKCδ positive neurons in the caudal CeA, suggesting a functional interaction between these cell types (Chou et al., 2022; Han et al., 2015; Bowen et al., 2023). Quantitative analysis of pERK cells within CGRPR and PKCδ neuronal populations revealed interesting proportions of exclusive and co-expressed cells. The presence of exclusive pERK cells in both CGRPR and PKCδ expressing neurons implies that these pathways may independently contribute to ERK activation in specific subpopulations of neurons. A noteworthy finding in our study was the substantial co-expression of pERK in neurons expressing both CGRPR and PKCδ in the CeA. This specify a potential functional convergence of these cells in the pain pathway, playing a critical role in pain modulation and sensitization processes within the CeA. A recent study on chronic migraine in a mouse model showed a significant increase in pERK expression in the CeA, and these pERK-positive neurons were predominantly composed of PKCδ positive neurons that also co-expressed CGRP receptors (Chou et al., 2022). These findings further strengthen the idea that CeA-CGRPR-expressing cells play a role in pain pathophysiology.

### Rostro-caudal variability in intrinsic excitability of CeA-CGRPR neurons in the context of neuropathic pain

Despite the well-established involvement of CGRP in pain within the CeA (Han et al., 2005, 2010), the intrinsic excitability of CGRPR-expressing CeA neurons has received limited attention in previous research. To bridge this knowledge gap, this study employs patch clamp electrophysiology to assess the intrinsic excitability of these neurons in the context of neuropathic pain. The findings reveal variations in the intrinsic excitability of CeA CGRPR neurons across different rostro-caudal levels, consistent with a previous study by Bowen et al. (2020). This show that the functional diversity of CGRPR neurons based on their rostro-caudal location is a general characteristic within the CeA and not specific to the neuropathic pain state. Furthermore, our study shows that these neurons display heightened excitability in response to higher current amplitudes, indicating their capacity for strong excitatory responses. However, the most noteworthy observation is the significant increase in excitability specifically at posterior rostro-caudal levels in mice with cuff implants. This finding implies that CGRPR-expressing neurons in the posterior CeA may play a more prominent role in processing nociceptive signals during neuropathic pain states. A recent study by Bowen et al. (2023) provided further support for our hypothesis regarding the heterogeneity of CeA-CGRPR neurons at different rostro-caudal levels. In their study, they demonstrated that rostral CeA and caudal CeA neurons mediate distinct functions through their complementary inputs and distinct outputs. This finding aligns with our observations of variations in intrinsic excitability of CeA-CGRPR neurons across different rostral-caudal levels in the context of neuropathic pain. The rostral-caudal differences observed here may also explain why no differences in intrinsic excitability of CeA neurons were found in a recent study between control mice and mice with a spared nerve injury in response to CGRP applied in slices (Trail et al., 2026). In that study, experiments were completed only in the posterior CeA.

Together, these studies emphasize the functional diversity of CeA-CGRPR neurons and shed light on the complex role they play in pain processing and modulation.

### CeA-CGRPR-expressing neurons are a key modulator of diverse pain modalities

Through pERK activation and *ex vivo* electrophysiology, we observed that CeA-CGRPR neurons become sensitized during pain states, indicating their significance in pain modulation. To establish a causal link between their activity and pain behavior, we employed a chemogenetic approach in a neuropathic mouse model, allowing us to selectively inhibit or activate CeA-CGRPR neurons. The precision of our chemogenetic targeting within the CeA was validated through the significant reduction or enhancement of Fos expression in transduced cells following chemogenetic inhibition or activation, respectively.

At the behavioral level, the inhibition of CeA-CGRPR neurons resulted in a significant reversal of cuff-induced hypersensitivity, highlighting their important role in mediating tactile and pressure hypersensitivity after nerve injury. Notably, the effects of inhibiting these neurons were specific to nerve injury, as there was no measurable impact in sham-treated mice or in the right hindpaw contralateral to the left nerve cuff. Furthermore, chemogenetic inhibition of CGRPR-positive neurons in the CeA also reduced hypersensitivity responses to cold and heat stimulation of the hindpaws, further underscoring their involvement in diverse pain modalities. The chemogenetic activation of these neurons induced significant hypersensitivity to various pain modalities, including tactile, pinch, cold, and heat stimulation. This observation emphasize the potent influence of these neurons in modulating pain responses and indicate that their aberrant activation may contribute to pain sensitization even without any external injury.

Behavioral results from our study are consistent with prior pharmacological experiments, emphasizing the key role of CeA-CGRP receptors in pain modulation. They demonstrated that administering CGRP1 receptor antagonists into the CeA effectively inhibits pain behaviors in arthritis and neuropathic pain models (Han et al., 2005, 2015; Preston, 2022). Furthermore, the localization of CGRP neuropeptide, which serves as a ligand for CGRP receptors, has been distinctly identified within the lateral and capsular domains of the CeA. The involvement of CGRP in regulating synaptic input originating from the PB to the CeA emphasizes the importance of CGRPR in synaptic plasticity within the CeA, with specific relevance to arthritis pain modulation (Han et al., 2005). Noteworthy, a recent pharmacological study with the spared nerve injury found that CGRP reduced cold hypersensitivity when CGRP was infused in the right CeA (left sciatic nerve injury) (Trail et al., 2026). The differences of our own study from that work may be related to methods (DREADD vs pharmacology), nerve injury model (cuff vs spared nerve), or timing of intervention. Overall, our findings further support the significance of CeA-CGRPR neurons in pain processing and contribute to the growing body of evidence suggesting that targeting these neurons may hold therapeutic potential for pain management.

### Sex-specific roles of CeA-CGRPR neurons in formalin-induced spontaneous pain behavior

Chronic pain affects a significant portion of the population, with females being disproportionately affected (Goldberg & McGee, 2011; Riley et al., 1998; Fillingim., et al 2009; Racine., et al 2012). However, many current analgesics are less effective in female patients due to their focus on male-specific pain processing mechanisms. Women often exhibit heightened sensitivity to various acute pain modalities and chronic pain conditions such as migraine, fibromyalgia, and irritable bowel disorder (Mogil, 2012; Unruh, 1996). While sex hormones play a role, recent research demonstrates that other biological factors, including genes, neurons, and immune cells (Beggs & Salter, 2010; Mogil, 2018, Mogil, 2020), operate differently in males and females regarding pain detection, transmission, and modulation.

Given the observed differences in pain experiences between men and women, understanding the sex-specific roles of CeA-CGRPR neurons in nociceptive behavior is critical. We found that chemogenetic inhibition of CeA-CGRPR neurons reduced cuff-induced hypersensitivity irrespective of sex. However, we observed sex-specific differences in the response to formalin-induced spontaneous pain, particularly in female mice. Notably, we discovered that chemogenetic inhibition of CeA-CGRPR neurons specifically reduced the second phase of formalin-induced pain responses in female mice but not in male mice. This observation highlights a sex-specific role of CeA-CGRPR neurons in modulating formalin-induced spontaneous pain, particularly at the phase associated with spinal sensitization. These results align with emerging evidence highlighting sex-specific differences in pain processing and modulation. Studies investigating the role of CGRP in pain modulation have also revealed sex-specific effects. For example, previous research has shown differential expression levels of CGRP-related genes in the spinal cord between males and females (Yadong et al., 2019). Moreover, CGRP-related pain behaviors, such as responses to neuropathic pain models (Peyton and Volker., 2022), often exhibit sex-specific patterns of sensitivity and responsiveness to CGRP receptor antagonists. CGRP signaling in the context of migraine has been a major therapeutic focus in the last decade. Evidence is emerging now for strong efficacy of CGRP inhibition in women compared to men (Porreca et al., 2024) matching up with existing preclinical data show sex-differences in CGRP signaling in animal models of migraine (Schou et al., 2017; Lewter et al., 2025).

While the precise mechanisms underlying these differences remain to be fully understood, our study contributes to a growing body of literature emphasizing the importance of considering sex as a biological variable in pain research. Further investigation into the molecular and circuit-level mechanisms governing the sex-specific roles of CeA-CGRPR neurons in pain modulation is warranted

## Acknowledgements

We would like to thank Dr. Richard Palmiter (University of Washington) for generously providing the CGRPR-cre mice and Dr. Maria L. Torruella-Suarez for providing feedback on histology data. pAAV-hSyn-DIO-hM3D(Gq)-mCherry (Addgene viral prep # 44361-AAV8; http://n2t.net/addgene:44361; RRID: Addgene_44361), pAAV-hSyn-DIO-hM4D(Gi)-mCherry (Addgene viral prep # 44362-AAV8; http://n2t.net/addgene:44362; RRID: Addgene_44362), and pAAV-hSyn-DIO-mCherry (Addgene viral prep # 50459-AAV8; http://n2t.net/addgene:50459; RRID: Addgene_50459) were a gift from Bryan Roth. We thank Phuoc Pham and George Dold (NIH Section on Instrumentation) for helping to design and fabricating the custom-built instruments used in this study. Figures were created with BioRender.com.

## Author Contribution

Conceptualization, YC, SS; Investigation, SS, AD, BN, SC, LAL, WF, JL, SV, TDW, JTR; Writing, SS, AD, BN, LAL, BJK, YC; Supervision, YC, BJK; Funding Acquisition, YC, SS, BJK.

---

**Supplementary Figure 1: Expression of CGRPR and PKCδ in the CeA is Consistent Between Sexes and Hemispheres.**

**(A).** Representative images of coronal sections containing the CeA expressing CGRPR tdTomato (red) and immunostained for PKCδ (cyan). Scale bars represent 1000 μm in the top row, 100 μm in the next three rows, and 10 μm in bottom row.

**(B).** Quantification of cells positive for CGRPR. Left panel – rostrocaudal level distribution showing average number of positive cells for levels -0.70 to -2.06 (n = 4 male mice, 3 female mice, 10 to 12 slices per mouse). Middle panel – average number of positive cells in entire CeA (n = 3 male mice, 3 female mice, 11 slices per mouse). Right panel – average number of positive cells per subnuclei region (n = 3 male mice, 3 female mice, 11 slices per mouse).

**(C).** Quantification of cells positive for PKCδ. Left panel – rostrocaudal level distribution showing average number of positive cells for levels -0.70 to -2.06 (n = 4 male mice, 3 female mice, 10 to 12 slices per mouse). Middle panel – average number of positive cells in entire CeA (n = 3 male mice, 3 female mice, 11 slices per mouse). Right panel – average number of positive cells per subnuclei region (n = 3 male mice, 3 female mice, 11 slices per mouse).

**(D).** Quantification of cells co-labeled for CGRPR and PKCδ. Left panel – rostrocaudual levels distribution showing average number of positive cells for levels -0.70 to -2.06 (n = 4 male mice, 3 female mice, 10 to 12 slices per mouse). Middle panel – average number of positive cells in entire CeA (n = 3 male mice, 3 female mice, 11 slices per mouse). Right panel – rostrocaudal level distribution of percentage of co-labeled cells, levels -0.70 to - 2.06 (n = 4 male mice, 3 female mice, 10 to 12 slices per mouse).

## References

Abbott FV, Franklin KBJ, Westbrook RF (1995) The formalin test: Scoring properties of the first and second phases of the pain response in rats. Pain 60:91–102.

Adke AP, Khan A, Ahn HS, Becker JJ, Wilson TD, Valdivia S, Sugimura YK, Gonzalez SM, Carrasquillo Y (2021) Cell type specificity of neuronal excitability and morphology in the central amygdala. eNeuro 8:ENEURO.0402-20.2020.

Ahrens S, Wu MV, Furlan A, Hwang GR, Paik R, Li H, Penzo MA, Tollkuhn J, Li B (2018) A central extended amygdala circuit that modulates anxiety. Journal of Neuroscience 38:5567–5583.

Allen HN, Chaudhry S, Hong VM, Lewter LA, Sinha GP, Carrasquillo Y, Taylor BK, Kolber BJ (2023) A parabrachial-to-amygdala circuit that determines hemispheric lateralization of somatosensory processing. Biological Psychiatry 93:370–381.

Beggs S, Salter MW (2010) Microglia–neuronal signalling in neuropathic pain hypersensitivity 2.0. Current Opinion in Neurobiology 20:474–480.

Benbouzid M, Pallage V, Rajalu M, Waltisperger E, Doridot S, Poisbeau P, Freund-Mercier MJ, Barrot M (2008) Sciatic nerve cuffing in mice: A model of sustained neuropathic pain. European Journal of Pain 12:591–599.

Bernard JF, Huang GF, Besson JM (1992) Nucleus centralis of the amygdala and the globus pallidus ventralis: Electrophysiological evidence for an involvement in pain processes. Journal of Neurophysiology 68:551–569.

Bowen AJ, Huang YW, Chen JY, Pauli JL, Campos CA, Palmiter RD (2023) Topographic representation of current and future threats in the mouse nociceptive amygdala. Nature Communications 14:196.

Calhoon GG, Tye KM (2015) Resolving the neural circuits of anxiety. Nature Neuroscience 18:1394–1404.

Carrasquillo Y, Gereau RW (2007) Activation of the extracellular signal-regulated kinase in the amygdala modulates pain perception. Journal of Neuroscience 27:1543–1551.

Carter ME, Soden ME, Zweifel LS, Palmiter RD (2013) Genetic identification of a neural circuit that suppresses appetite. Nature 503:111–114.

Choi Y, Yoon YW, Na HS, Kim SH, Chung JM (1994) Behavioral signs of ongoing pain and cold allodynia in a rat model of neuropathic pain. Pain 59:369–376.

Chou TM, Lee ZF, Wang SJ, Lien CC, Chen SP (2022) CGRP-dependent sensitization of PKCδ-positive neurons in central amygdala mediates chronic migraine. The Journal of Headache and Pain 23:157.

Colburn RW, Lubin ML, Stone DJ Jr, Wang Y, Lawrence D, Brandt MR, Liu Y, Flores CM, Qin N (2007) Attenuated cold sensitivity in TRPM8 null mice. Neuron 54:379–386.

D’Hanis W, Linke R, Yilmazer-Hanke DM, Schwegler H (2007) Topography of CGRP-containing neurons in the rat parabrachial nucleus projecting to the amygdala. Neuroscience Letters 412:146–150.

Dai Y, Zhang Y, Zhao J, McDonald JW (2002) Differential activation of MAPK in injured and uninjured dorsal root ganglion neurons following chronic constriction injury of the sciatic nerve in rats. European Journal of Neuroscience 20:2881–2895.

Dobolyi A, Irwin S, Makara G, Usdin TB, Palkovits M (2005) Calcitonin gene-related peptide-containing pathways in the rat forebrain. Journal of Comparative Neurology 489:92–119.

Ehrlich I, Humeau Y, Grenier F, Ciocchi S, Herry C, Lüthi A (2009) Amygdala inhibitory circuits and the control of fear memory. Neuron 62:757–771.

Fadok JP, Markovic M, Tovote P, Lüthi A (2018) New perspectives on central amygdala function. Current Opinion in Neurobiology 49:141–147.

Fillingim RB, King CD, Ribeiro-Dasilva MC, Rahim-Williams B, Riley JL III (2009) Sex, gender, and pain: A review of recent clinical and experimental findings. The Journal of Pain 10:447–485.

Goldberg DS, McGee SJ (2011) Pain as a global public health priority. BMC Public Health 11:770.

Han JS, Li W, Neugebauer V (2005) Critical role of calcitonin gene-related peptide 1 receptors in the amygdala in synaptic plasticity and pain behavior. Journal of Neuroscience 25:10717–10728.

Han JS, Adwanikar H, Li Z, Ji G, Neugebauer V (2010) Facilitation of synaptic transmission and pain responses by CGRP in the amygdala of normal rats. Molecular Pain 6:10.

Han S, Soleiman MT, Soden ME, Zweifel LS, Palmiter RD (2015) Elucidating an affective pain circuit that creates a threat memory. Cell 162:363–374.

Hargreaves K, Dubner R, Brown F, Flores C, Joris J (1988) A new and sensitive method for measuring thermal nociception in cutaneous hyperalgesia. Pain 32:77–88.

Hurst JL, West RS (2010) Taming anxiety in laboratory mice. Nature Methods 7:825–826.

Iyengar S, Ossipov MH, Johnson KW (2017) The role of calcitonin gene-related peptide in peripheral and central pain mechanisms including migraine. Pain 158:543–559.

Janak PH, Tye KM (2015) From circuits to behaviour in the amygdala. Nature 517:284–292.

Ji Y, Rizk A, Voulalas P, Aljohani H, Akerman S, Dussor G, Keller A, Masri R (2019) Sex differences in the expression of calcitonin gene-related peptide receptor components in the spinal trigeminal nucleus. Neurobiology of Pain 6:100031.

Lewter LA, Arnold RL, Narosov NB, Dussor G, Kolber BJ (2025) Sex differences in the effects of calcitonin gene-related peptide signaling on migraine-like behavior in animal models: A narrative review. Frontiers in Neurology 16:1603758.

Li H, Penzo MA, Taniguchi H, Kopec CD, Huang ZJ, Li B (2013) Experience-dependent modification of a central amygdala fear circuit. Nature Neuroscience 16:332–339.

Moscarello JM, Penzo MA (2022) The central nucleus of the amygdala and the construction of defensive modes across the threat-imminence continuum. Nature Neuroscience 25:999–1008.

Mogil JS (2012) Sex differences in pain and pain inhibition: Multiple explanations of a controversial phenomenon. Nature Reviews Neuroscience 13:859–866.

Mogil JS (2018) Sex-based divergence of mechanisms underlying pain and pain inhibition. Current Opinion in Behavioral Sciences 23:113–117.

Mogil JS (2020) Qualitative sex differences in pain processing: Emerging evidence of a biased literature. Nature Reviews Neuroscience 21:353–365.

Neugebauer V, Li W, Bird GC, Bhave G, Gereau RW (2003) Synaptic plasticity in the amygdala in a model of arthritic pain: Differential roles of metabotropic glutamate receptors 1 and 5. Journal of Neuroscience 23:52–63.

Neugebauer V (2022) Amygdala pain mechanisms. In: Handbook of Clinical Neurology, Vol 184, pp 203–223.

O’Leary TP, Kendrick RM, Bristow BN, Sullivan KE, Wang L, Clements J, Lemire AL, Cembrowski MS (2022) Neuronal cell types, projections, and spatial organization of the central amygdala. iScience 25:105497.

Paxinos G, Franklin KBJ (2001) *The mouse brain in stereotaxic coordinates*, 2nd Ed. San Diego: Academic Press.

Penzo MA, Robert V, Tucciarone J, De Bundel D, Wang M, Van Aelst L, Darvas M, Parada LF, Palmiter RD, He M, Huang ZJ, Li B (2015) The paraventricular thalamus controls a central amygdala fear circuit. Nature 519:455–459.

Porreca F, Navratilova E, Hirman J, van den Brink AM, Lipton RB, Dodick DW (2024) Evaluation of outcomes of calcitonin gene-related peptide-targeting therapies for acute and preventive migraine treatment based on patient sex. Cephalalgia 44.

Presto P, Neugebauer V (2022) Sex differences in CGRP regulation and function in the amygdala in a rat model of neuropathic pain. Frontiers in Molecular Neuroscience 15:928587.

Racine M, Tousignant-Laflamme Y, Kloda LA, Dion D, Dupuis G, Choinière M (2012) A systematic review of sex/gender differences in pain perception. Pain 153:602–618.

Randall LO, Selitto JJ (1957) A method for measurement of analgesic activity on inflamed tissue. Archives Internationales de Pharmacodynamie et de Thérapie 111:409–419.

Riley JL III, Robinson ME, Wise EA, Myers CD, Fillingim RB (1998) Sex differences in the perception of noxious experimental stimuli: A meta-analysis. Pain 74:181–187.

Russo AF (2015) Calcitonin gene-related peptide (CGRP): A new target for migraine. Annual Review of Pharmacology and Toxicology 55:533–552.

Schou WS, Ashina S, Amin FM, Goadsby PJ, Ashina M (2017) Calcitonin gene-related peptide and pain: A systematic review. The Journal of Headache and Pain 18:34.

Shinohara Y, Saito Y, Kumamoto E, Shibata M (2017) Calcitonin gene-related peptide facilitates synaptic transmission and pain responses in the amygdala via the PKC pathway. Neuroscience 346:180–190.

Singh S, Wilson TD, Valdivia S, Benowitz B, Chaudhry S, Ma J, Adke AP, Soler-Cedeño O, Velasquez D, Penzo MA, Carrasquillo Y (2022) An inhibitory circuit from central amygdala to zona incerta drives pain-related behaviors in mice. eLife 11:e68760.

Trail AD, Allen HN, Paul B, Nelson TS, Widner JA, Lewter L, Tack SH, Neilan RM, Kolber BJ (2026) Amygdalar calcitonin gene-related peptide-driven effects of cold sensitivity induced by peripheral neuropathy in mice. Journal of Pain 41:106199.

Unruh AM (1996) Gender variations in clinical pain experience. Pain 65:123–167.

Veinante P, Yalcin I, Barrot M (2013) The amygdala between sensation and affect: A role in pain. Journal of Molecular Psychiatry 1:9.

Wilson TD, Valdivia S, Khan A, Ahn HS, Adke AP, Martinez Gonzalez S, Sugimura YK, Carrasquillo Y, Palmiter RD (2019) Dual and opposing functions of the central amygdala in the modulation of pain. Cell Reports 29:3324–3336.

Xu W, Lundeberg T, Wang YT, Li Y, Yu LC (2003) Antinociceptive effect of calcitonin gene-related peptide in the central nucleus of amygdala: Activating opioid receptors through the amygdala–periaqueductal gray pathway. Neuroscience 118:1015–1022.

Zhuang ZY, Gerner P, Woolf CJ, Ji RR (2005) ERK is sequentially activated in neurons, microglia, and astrocytes by spinal nerve ligation and contributes to mechanical allodynia. Pain 114:149–159.

