## Supplementary figures and images for "CGRP receptor-expressing neurons in the central amygdala contributes to injury-induced pain hypersensitivity"

### Supplemental figure

#### Supplementary Figure 1

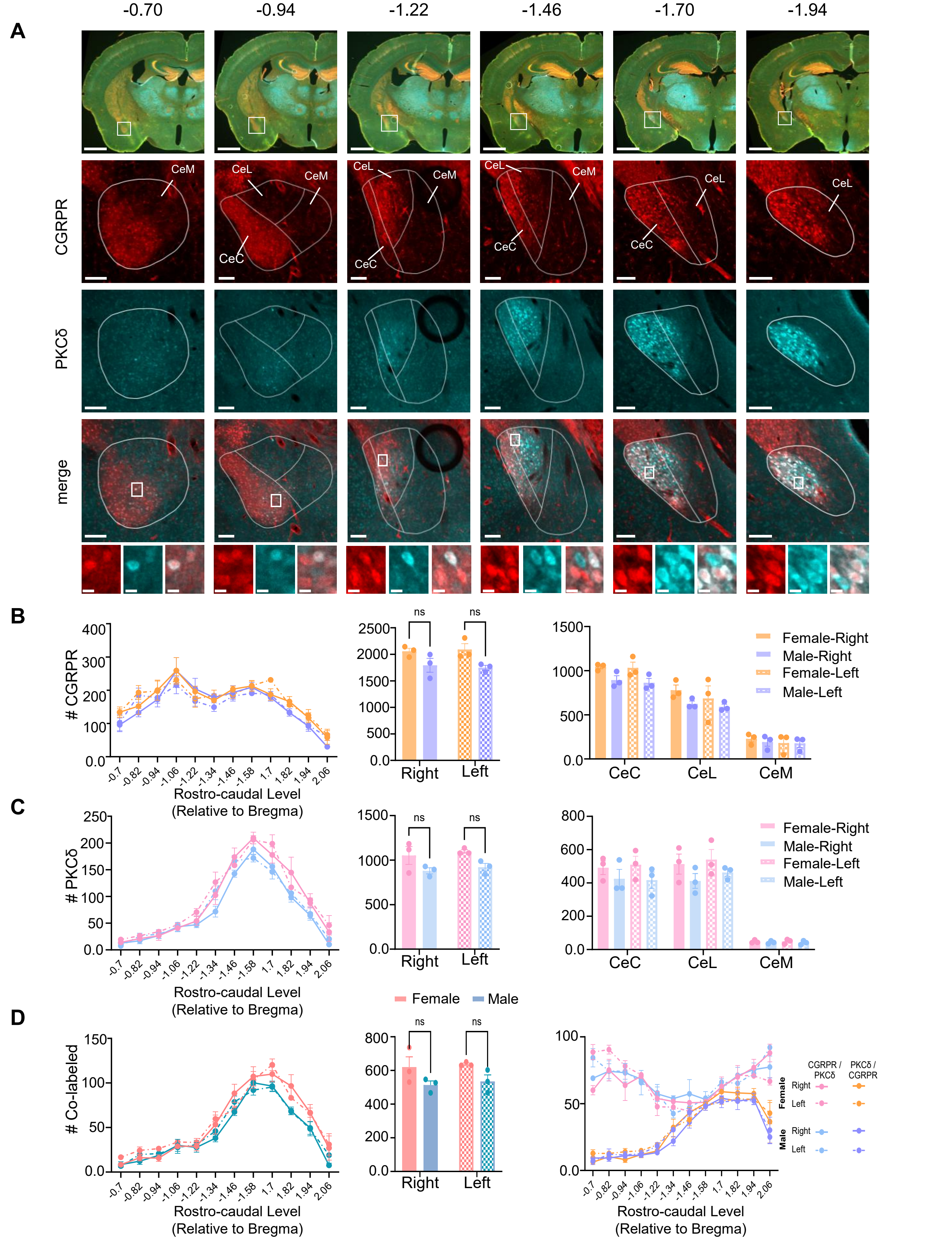
